# T Cell Receptors for Antigen on Intraepitheial Cytolytic T Lymphocytes in Celiac Disease Engage Enterocyte HLA-E and HLA-B

**DOI:** 10.1101/2024.09.03.610908

**Authors:** JE Johnson, K Agrawal, RS Al-Lamki, F Zhang, Wang Xi, Z Tobiasova, SA Taleb, S Liburd, L Rodriguez, AJ Martins, RA Flavell, ME Robert, E Sefik, JS Pober

## Abstract

We compared duodenal biopsies showing active celiac disease (CeD) to normal controls using single cell RNA sequencing, cyclic immunofluorescence, RNAScope and proximity ligation assays. There is increased infiltration of villous but not crypt epithelium T cells bearing either αβ or γδ T cell receptors (TCRs) in CeD. Both T cell subsets are activated cytotoxic T lymphocytes (CTLs) and surprisingly are the predominant mucosal source of IFNγ. In response to this IFNγ,villous but not crypt enterocytes show an IFNγ signature, including nuclear phospho-STAT1 protein, class II HLA molecules and IFNγ-inducible chemokines known to recruit CTLs (e.g., CCL3, CCL4, CXCL10, and CXCL11) and receptors for these chemokines are expressed on the infiltrating CTLs. Villous enterocytes also display increased HLA-E and HLA-B mRNAs and proteins. Bioinformatic analyses (NICHES) and proximity ligation assays show frequent binding of both αβ and γδ TCRs with enterocyte HLA-E or HLA-B, but not HLA-DR. In contrast, NKG2C, proposed as an alternative trigger of CTL activation, is infrequently-expressed and shows few interactions with HLA-E. Our data suggest that activated intraepithelial CTLs produce IFNγ which recruits additional CTLs and increases antigen-dependent killing of villous epithelium using either conventional or HLA-E antigen presentation.

**Signifance:** Analyses of diagnostic biopsies from celiac disease patients reveals critical roles for IFNγ actions on enterocytes and a surprising role of HLA-E in antigen presntation. Activated cytotoxic T lymphocytes (CTLs) that are the primary source of mucosal IFNγ, leading to IFNγ signaling in villous enterocytes, including expression of phospho-STAT1 protein, class II HLA molecules, and chemokines that attract CTLs. Bioinformatic ligand receoptor analyses and proximity ligation assays showed frequent interactions between intraepithelial CTLs and enterocyte HLA-E or HLA-B, indicating that IFNγ production by the CTLs likely promotes further CTL recruitment and antigen-dependent killing of the villous epithelium.

## Introduction

Celiac disease (CeD) is the most common autoimmune disorder in the U.S., affecting at least 1% of the population. Ingestion of grains (wheat, barley or rye) containing gluten proteins (gliadins or glutenins) triggers an adaptive immune response that causes both gastrointestinal and systemic symptoms (1, 2). Lack of approved therapy other than dietary avoidance of gluten, which is difficult and often unsuccessful, highlights the need to identify new mechanisms with insights for developing therapeutics (3). Although serological titers of IgA antibodies to transglutaminase 2 (also known as tissue transglutaminase or TTG) or endomysium can aid in diagnosis, definitive diagnosis involves a duodenal mucosal biopsy showing histologic changes of increased ratios of T cells to enterocytes within the villous epithelium, villous flattening, and crypt elongation (4, 5).

CeD pathogenesis is multifactorial with genetics playing a key role. Almost all patients with CeD have inherited at least one copy of an HLA-DQ2 or DQ8 allele, normally found at a frequency of 30% in the total population (6–8). The presence of HLA-DQ2/DQ8 alleles provides a strong risk factor but is not sufficient in causing disease. Certain class I HLA alleles also confer a risk, although this may reflect linkage disequilibrium to HLA-DQ2 (6). DQ2/DQ8 alleles are enriched because the proteins they encode can uniquely bind certain gluten peptides that have been deaminated by TTG (9, 10). These deamidated gluten peptides (DGPs) are presented to CD4 T cells with relevant αβ T cell receptors for antigen (TCRs). Upon gluten challenge to an individual with established CeD, clonally expanded, DGP-reactive CD4 T cells are found both in the circulation and within the lamina propria (LP) of the small intestinal mucosa. In vitro, these CD4 T cells will express both IFN-g and IL-21 in response to presentation of DGP antigens (11–13) and have a phenotype that matches that of T peripheral helper cells found in multiple other autoimmune disorders (14, 15). DGP-reactive CD4 T cells likely provide help to B cells expressing surface immunoglobulins that recognize DGPs or TTG/DGP complexes. These B cells give rise to plasma cells that produce IgG and IgA antibodies against DGPs or TTG, many of which also localize to the LP of the small intestine. CD4 T cell-derived IL-21 may also help naïve CD8 T cells to differentiate into cytotoxic T lymphocyte (CTL) effector cells (11, 16).

Lymphocyte recruitment to the duodenum is likely driven by chemokines. The homeostatic chemokine CCL25 is constitutively produced by the small intestine and contributes to homing of CCR9 expressing CTLs, CD4 T cells, and other cell types to the intestinal LP (17). In CeD activated CTLs further home to the epithelial lining of the small intestinal villi (18). Inflammatory chemokines in addition to tissue-specific chemokines, particularly those that bind to chemokine receptor CCR5 (i.e., CCL3, 4 or 5) and/or to CXCR3 (i.e., CXCL9, 10, or 11), likely contribute to this recruitment (19, 20). Although prior studies have shown an increase in several of these chemokines in CeD, in particular those that bind to CXCR3, less is known about expression of these molecules in different tissue compartments and in specific recruitment of CTLs to the villous epithelium compared to the crypt epithelium or LP stroma (21, 22).

Infiltrating CTLs within the villous epithelium, which replace the resident intraepithelial T cell populations (23), are presumed to kill enterocytes, leading to villous flattening.CTL killing may by mediated either by granule exocytosis of secreted cytotoxic molecules (e.g., perforin, granzyme B, or granulysin) or by inducing caspase activation on target cells through engagement of “death receptors” on the surface (e.g Fas by Fas ligand on T cells) (24). Both pathways of CTL killing are contact-dependent, requiring recognition of a ligand on the target cell by an activating receptor on the CTL in order to trigger killing (25). Generally, this involves TCR signaling initiated by recognition of cognate peptide bound to a class I HLA molecule expressed on the target cell. However, the triggering of intraepithelial CTLs in CeD is a matter of controversy. When a CeD patient is challenged with gluten, expanded CTL clones bearing either αβ or γδ TCRs increase within the bloodstream and the same TCRs may be found in the villous lining of the duodenum, suggestive of an antigen-specific response to gluten similar to that seen for CD4 T cells (26–28). Further, some CD8 T cell responses to gluten peptides being presented in association with class I HLA-A or B alleles have been reported, including CD8 cells within the mucosa (29–31). However, there has been no direct demonstration of gluten peptide presentation by enterocytes and TCR sequencing has identified the whole intraepithelial T lymphocyte population to be highly polyclonal, suggesting independence of antigen (32–34). Jabri and colleagues have proposed a broadly accepted alternative hypothesis in which CTLs may bypass TCR-mediated activation and instead, use activating NK cell receptor CD94/NKG2C that binds to HLA-E independently of peptide recognition (35). NKG2C can be induced on CTL by high concentrations of IL-15 and, by signaling through adaptor protein DAP12, can trigger granule exocytosis and killing (35, 36). This view is supported by evidence of elevated expression of IL-15 and of HLA-E by enterocytes in CeD (35, 37). It is important to note that HLA-E, in addition to its role as a ligand for NK cell receptors, can also be a functional antigen presenting element for a few known peptides associated with significant human pathogens (38).

Although active CeD is widely appreciated to be associated with elevated levels of IFNγ, the sources and specific roles played by this cytokine in CeD are not well defined. Our new spatial data suggest that intraepithelial CTL and not CD4 T cells in the LP, are the primary and perhaps sole source of IFN-γ in CeD. Further, we present evidence that two key actions of IFN-γ on villous enterocytes are to induce inflammatory chemokines that recruit CTL and increase expression of MHC molecular targets, especially HLA-E, for triggering CTL activation and killing. By computational analysis with NICHES and direct examination of protein-protein interactions with proximity ligation assays, we find that intraepithelial CTL recognize HLA-E principally through their TCRs rather than through NKG2C. We also find significant levels of interactions of TCRs with conventional class I HLA-B molecules. These new findings suggest that intraepithelial CTL likely recognize gluten peptide antigens presented by enterocytes, which present antigen using both HLA-B and HLA-E

## Results

### Transcriptional landscape of CeD at single cell level

We began our analysis of CeD using single cell RNA sequencing (scRNA-seq) to gain a non-biased view of the transcriptional landscape of the duodenum. We analyzed fresh research biopsies from 4 patients with active CeD, confirmed by diagnostic biopsies from the same endoscopy, as well as 2 healthy control biopsies without specific pathological changes. Our analysis captured every major intestinal mucosal cell type including all subsets of immune cells, epithelial cells and stromal cells (Fig. 1a) (39). Among these, enterocytes and activated T cells exhibited the most significant differences between CeD biopsies and controls (Fig 1a-c). We have not been able to cleanly separate T cells from the epithelial layer and the LP despite initially enriching for epithelium in our cell suspension procedure. While CD103 is sometimes used to identify intraepithelial T cells, its also present in circulating CTL in CeD patients and not restricted to the epithelial layer in mucosal biopsies (27). In the initial UMAP embedding, CD8+ TCR αβ and γδ cells could not be separated due to their similar transcriptomes. To address this, we focused on T cell subsets and increased cluster resolution (Fig. 1b, c). Both TCR αβ and TCR γδ T cell subsets in CeD were enriched for markers of T cell activation and cytotoxicity and clustered distinctly compared to their counterparts in control biopsies. (Fig. 1b, c).

**Figure 1.**
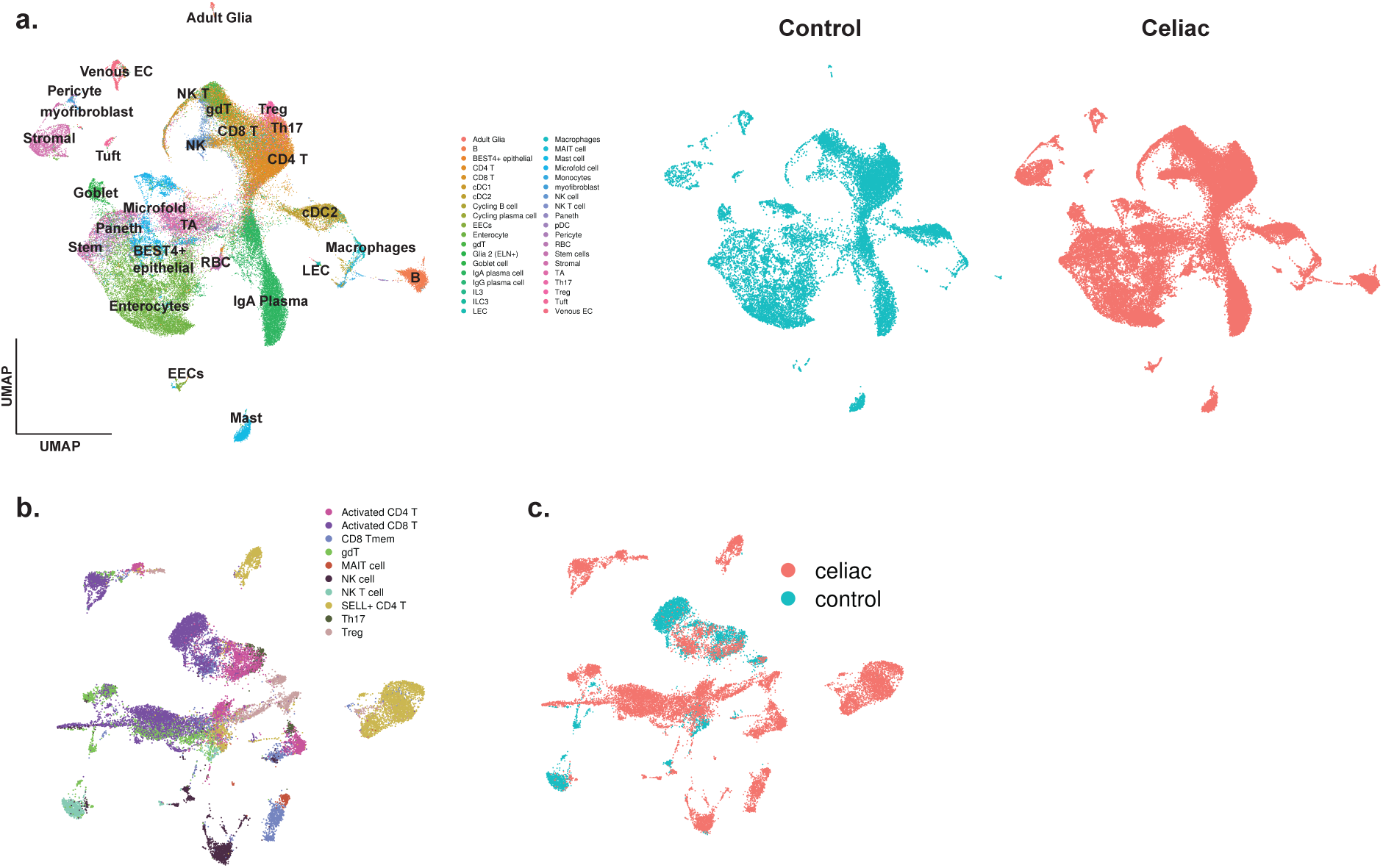
scRNA-seq analysis of CeD and control biopsies. **(a).** Left: UMAP embedding and annotation of three celiac biopsies and two control biopsies. Annotation was performed using Celltypist and the human gut reference atlas. Right: Visualization of control and celiac group cells in the same UMAP embedding. **(b).** UMAP embedding of T and NK cell clusters and annotation of T cell subpopulations. **(c).** UMAP embedding showing distribution of cells from celiac vs control groups

### Characteristic Cellular Composition of Villous/Surface Epithelium in CeD and Control Biopsies

Informed by the scRNA-seq data, we applied spatial approaches to focus further on the interactions between T cells and enterocytes within the epithelial lining. Unlike analyses of isolated cells, spatial analyses allow for clear distinction between the epithelial and LP compartments. We first applied immunofluorescence microscopy (IF) to formalin-fixed paraffin-embedded (FFPE) diagnostic duodenal biopsies from normal controls and from CeD patients with severe villous flattening (Marsh grade 3B and C). Enterocytes were initially identified using pan-enterocyte marker E-cadherin (Fig. 2a). The villous epithelial compartment, whose boundaries can be difficult to define in CeD, was distinguished from crypt epithelium by ApoA4 expression and by staining of the transitional zone between villi and crypts with Ki-67 (Fig. 2b) (40). These epithelial markers along with general T cell marker CD3 allowed for accurate identification and quantification of the intraepithelial lymphocyte (IEL) population within the villous compartment compared to crypt and LP lymphocytes. Further analysis of these CD3 IEL populations revealed the majority of them to be CD8 positive (Fig. 2c). No CD4 T cells were detected within the epithelial lining (negative data not shown). In accordance with established diagnostic practice (41, 42), we determined the number of T cells per 100 epithelial cell nuclei (identified by DAPI staining). Diagnostic criteria for villous epithelial lymphocytosis in CeD include a threshold >25 intraepithelial lymphocytes per 100 epithelial cells (43). Our samples of CeD biopsies have an average of 36.1 ± 2.18 (SEM) CD3 T cells per 100 villous nuclei, and our normal control biopsies have a mean of 14.5 ± 2.97 CD3 T cells per 100 villous nuclei (Fig. 2d). These findings are consistent with prior reports of increased T cell numbers in the duodenal mucosa in active CeD (41, 42). Further breakdown of these IELs based on TCR staining showed increases in both TCR αβ and γδ populations, with TCR αβ populations expanding the most compared to controls (Fig. 2e, f). In contrast to the increased ratios of T cells to enterocytes within the villi in CeD, T cell numbers in the crypts of CeD biopsies showed no statistical difference in total and TCR αβ T cell ratios to enterocytes vs. controls and TCR γδ cells in the crypt were only slightly increased compared to controls (Fig. 2g-i). These observations indicate that our biopsy specimens are concordant with those used in prior studies.

**Figure 2.**
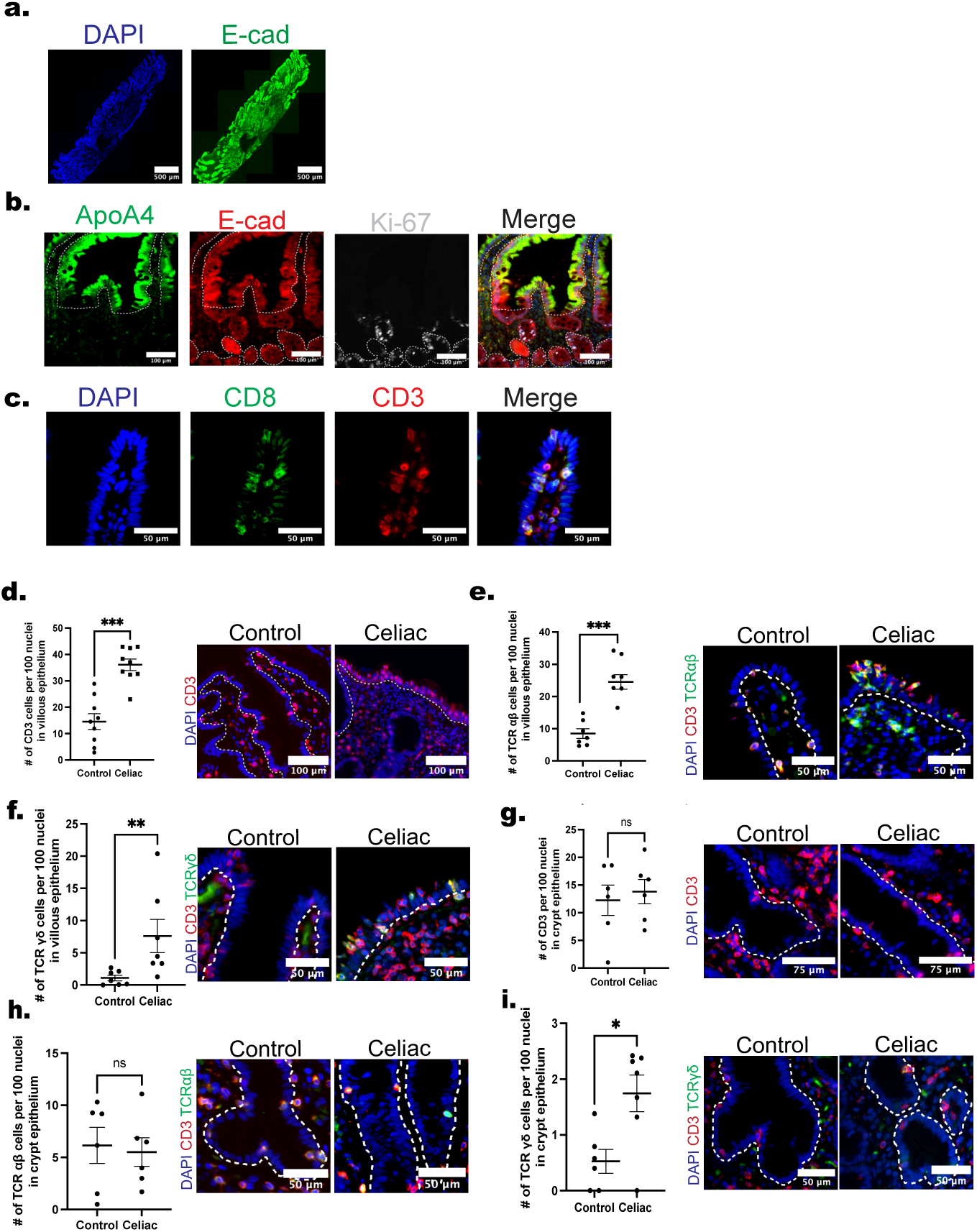
Quantitation of total CD3, TCRαβ and TCRγδ T cells within the duodenal villous and crypt epithelium in CeD vs. control biopsies. **(a).** Duodenal tissue was stained with pan-epithelial cell marker E-cadherin (green) and DAPI (blue) to assess tissue morphology. (**b**). ApoA4 (green) and Ki-67 (white) markers identify the villous and crypt epithelium respectively. The dotted white lines follow the basement membrane that separates villous and crypt epithelium from the LP. **(c).** Representative image of CD3 cells (red) in villus co-staining with CD8 (green). **(d).** Quantitation of the frequency villous epithelium infiltration of total CD3 T cells (N=9), (**e).** TCRαβ (N=7), **(f)** and TCRγδ T cell (N = 7)**. (g).** Quantitation of the frequency of crypt epithelium infiltration of total CD3 T cells (N=6), (**h).** TCRαβ (N=6), **(i)** and TCR γδ T cells (N = 6). T cells in representative images are labeled with CD3 (red) and TCRαβ or γδ (green). Values are presented as the ratio of T cells per enterocyte nuclei (stained with DAPI). Significance determined by t-test. \**P* ≤ 0.05; \*\**P* ≤ 0.01; \*\*\*\**P* ≤ 0.001.

### Characterization of intra-epithelial TCR αβ and TCR γδ T cell populations in CeD as CTLs

Going back to our transcriptional analysis, we analyzed subsets and transcriptional signatures associated with activated TCR αβ and TCR γδ T cell subsets in CeD, comparing them to what has been previously described. Among CD4+ TCR αβ cells, there was a marked reduction in IL-17 expressing T cells and an increase in regulatory T cells, which is consistent with other studies focusing on the roles of CD4+ T cells in celiac disease (28) (Fig. 3a). The same study also reported a population of innate-like T cells that expressed DAP12, CD247, and SH2D1B. A larger sample size may be needed to detect this rare subset as we could not detect this population in our analysis. Interestingly, CD4+ TCR αβ cells isolated from the gut expressed very little *IFNG* transcript(Fig. 3b) despite the abundant production of this cytokine when cells were stimulated in vitro (11–13). By contrast, activated CD8+ TCR αβ had elevated levels of *IFNG* (encoding IFNγ) as well as *GZMB* (encoding granzyme B) and *NKG7* (encoding natural killer cell granular protein 7) transcripts. CD8+TCR αβ and TCR γδ T cells were the two major cell types that expressed *IFNG* and both exhibited significant expression of *IFNG* transcripts, a feature that was largely absent in the control biopsies (Fig. 3b, c, Supplementary Fig. 1). Moreover, enterocytes that express villous defining marker APOA4+ showed elevated levels of *IFNGR1* (Fig. 3d, e), suggesting an increased responsiveness to IFNγ produced by CeD T cells. Our findings challenging most of earlier findings in CeD literature are consistent with two new studies, which also identified a similar population of IFNG expressing CD8+ T cells (28, 44).

**Figure 3.**
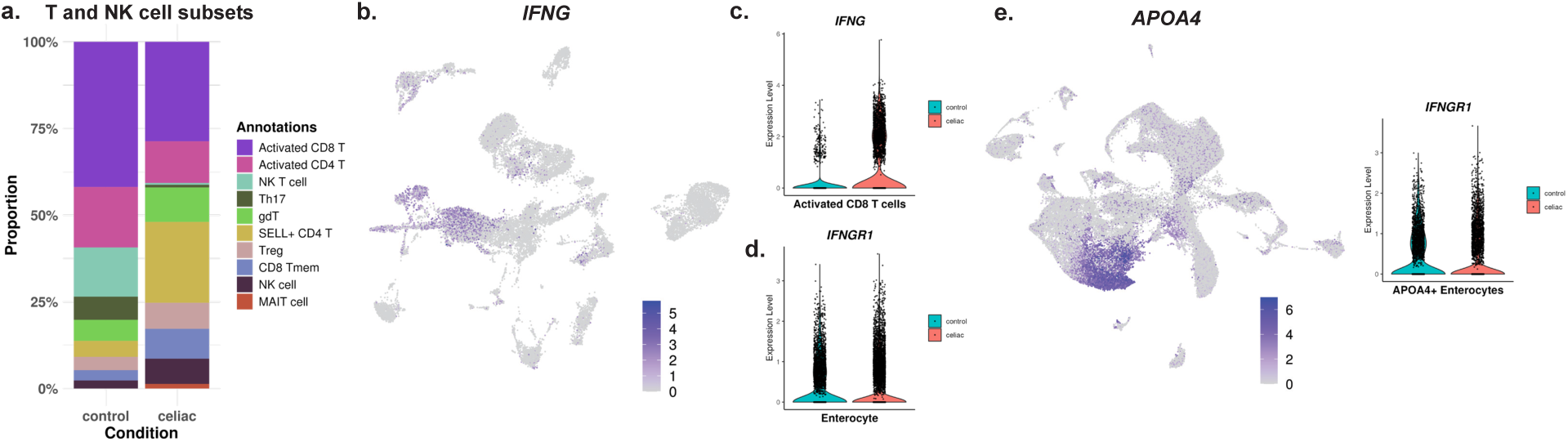
CD8+ T cells are the primary producers of *IFNG,* which is sensed by APOA4+ enterocytes. **(a).** Proportions of T cells and NK cells in control vs celiac groups **(b).** *IFNG* expression shown in UMAP embedding of total T and NK cell clusters **(c).** *IFNG* expression in activated CD8+ T cells in control vs celiac groups. *IFNG* expression across other T cell subsets including TCRγδ T cells are shown in Fig. S1**(d).** IFNGR1 expression in enterocytes in control vs celiac groups **(e).** IFNGR1 expression in villous enterocytes marked byAPOA4 expression in control vs celiac groups, displayed as violin plots.

Although transcriptional analysis clearly suggested accumulation of CTLs in CeD, our scRNA-seq experiments could not determine with certainty whether these cells were localized within the epithelial layer of the villi. Using IF, we found that a much greater percentage of intraepithelial T cells in CeD than controls express CD45RO, indicative of prior exposure to their cognate antigen (Fig. 4a,b). Mature CTL can also be distinguished by expression of cytosolic granules containing Granzyme B (GrzB) and by IFNγ production. We found 38.8 ±4.8% of villous intraepithelial T cells to be GrzB+ in CeD compared to 7.2 ±1.8% in controls (Fig. 4c, d). Increased GrzB expression was observed in both TCR αβ and γδ subsets in the villous (Supplementary Fig. 2a-b). Consistent with our scRNAseq analysis of isolated CD8 T cells., *IFNG* mRNA was found to be markedly elevated in villous epithelial T cells in CeD vs, control biopsies as assessed by RNAScope, but villous T cells in control biopsies did show some limited expression. (Fig. 4e-g). *IFNG* mRNA was lower in crypt intraepithelial T cells and essentially absent in T cells within the LP (Supplementary Fig. 3). Staining for T cell associated IFNγ protein was generally unscuccessful in both CeD and control biopsies (negative data not shown), consistent with secretion being concomitant with synthesis.

**Figure 4.**
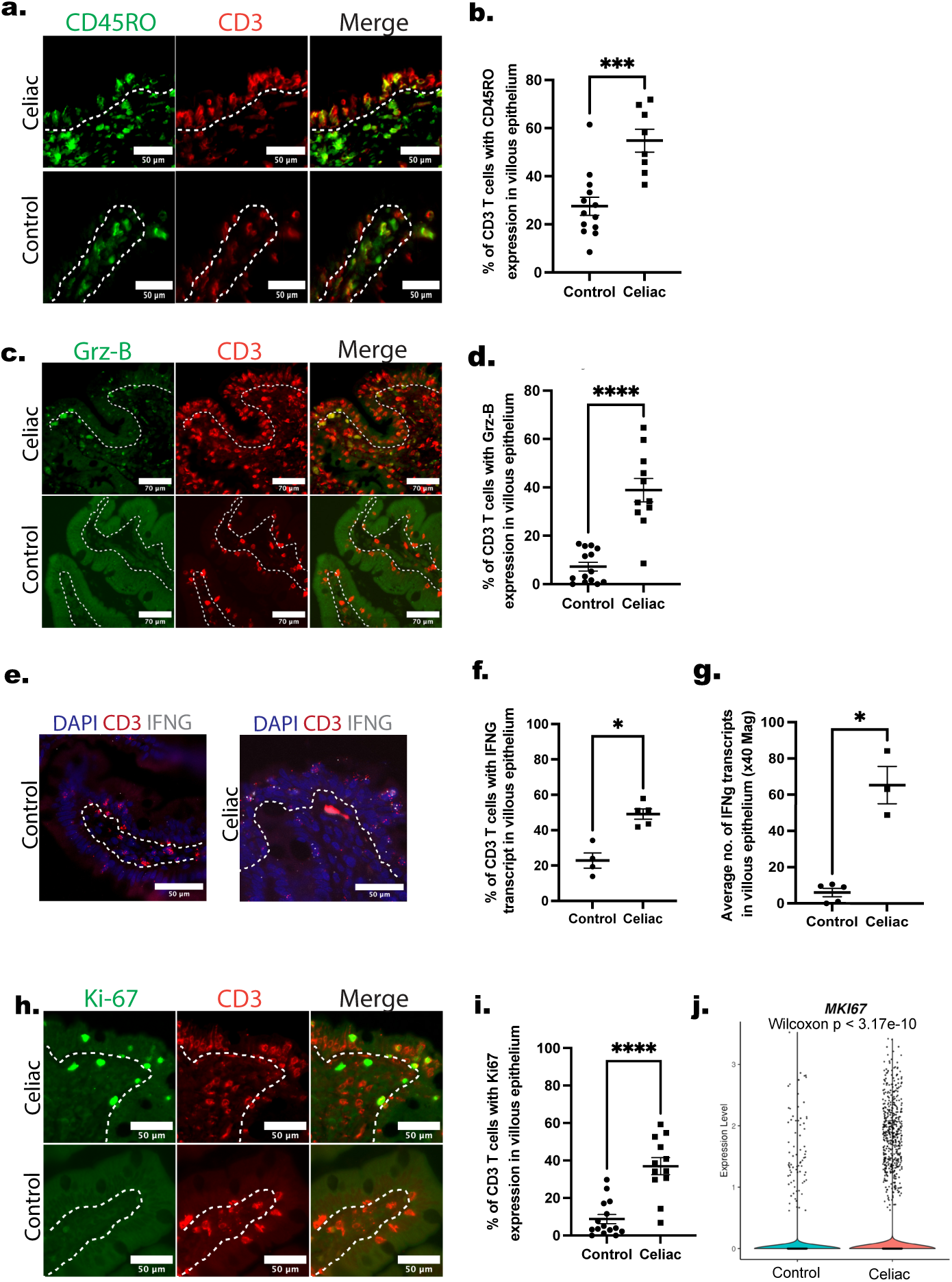
Identification of intraepithelial CD3 T cells in CeD as activated CTL. **(a, b).** CD45RO (N = 8) and **(c,d).** Granzyme B (N=11) protein expression was assessed by IF staining as the percentage of CD3 cells (red) positive for target protein within the villous epithelium of CeD vs control biopsies **(e).** Representative images of *IFNG* (white) and CD3 (red) transcript expression using RNAScope in control and celiac biopsies **(f).** Quantification of percent of CD3 T cells in villus with *IFNG* transcript. **(g).** Quantification of the number of *IFNG* transcripts per 40x image in the villous epithelium **(h,i)**. Ki-67 protein expression (green) by IF as the percentage of CD3 cells positive within the villous epithelium of biopsies (N=12) **(j).** Violin plots displaying expression values of MK167 in T cells from scRNA-seq. Significance determined by t-test. \**P* ≤ 0.05; \*\**P* ≤ 0.01; \*\*\*\**P* ≤ 0.001.

The synthesis of IFNγ, detected by *IFNG* transcript, not only indicates the potential function of the intraepithelial T cells as CTLs, but further suggests that these cells are in an activated state, typically as a result of antigen stimulation. CeD IELs were also found to be proliferative, with a significant increase in the percentage of intraepithelial T cells in CeD compared to controls expressing nuclear Ki-67 (Fig. 4h,i). Both TCR αβ and TCR γδ T cell subsets showed a similar increase in Ki-67 staining (Supplementary Fig. 2b). These findings are consistent with our scRNA-seq dataset that also shows a cycling population of T cells marked by MKI67 expression present at higher frequencies in CeD patients (Fig. 4j). Collectively, these data identify T cells within the residual villous epithelium as activated CTL and, contrary to prior interpretations, identify these cells as the overwhelmingly predominant producers of IFNγ within the duodenal mucosa of biopsies from active CeD patients, and largely confines IFNγ production to the residual villous epithelial lining.

### Evidence of IFNγ signaling in villous enterocytes in CeD

The presence of elevated *IFNG* within intraepithelial CTL in CeD led us to examine villous enterocyte populations for evidence of an IFNγ signaling response. The initial response to this cytokine involves Janus kinase (JAK)-mediated phosphorylation of tyrosine residues in the signal transducing and transcriptional activator 1 (STAT1) protein, followed by translocation of pSTAT1 to the nucleus (45). We found significantly increased levels of nuclear pSTAT1 in the villous enterocytes of CeD biopsies, with 59.1 ±6.5% of enterocyte nuclei pSTAT1+ compared to only 1.4 ±0.45% in normal controls (Fig. 5a, b). Phosphorylated STAT1 is also somewhat elevated in crypt enterocytes in CeD compared to controls, but this frequency is much lower than in villous enterocytes (Fig. 5c), consistent with fewer T cells and a reduced level of IFNγ in this compartment. Crypts contain a more varied population of enterocytes than villi and these observations do not rule out an IFNγ effect on some epithelial cell types within the crypts.

**Figure 5.**
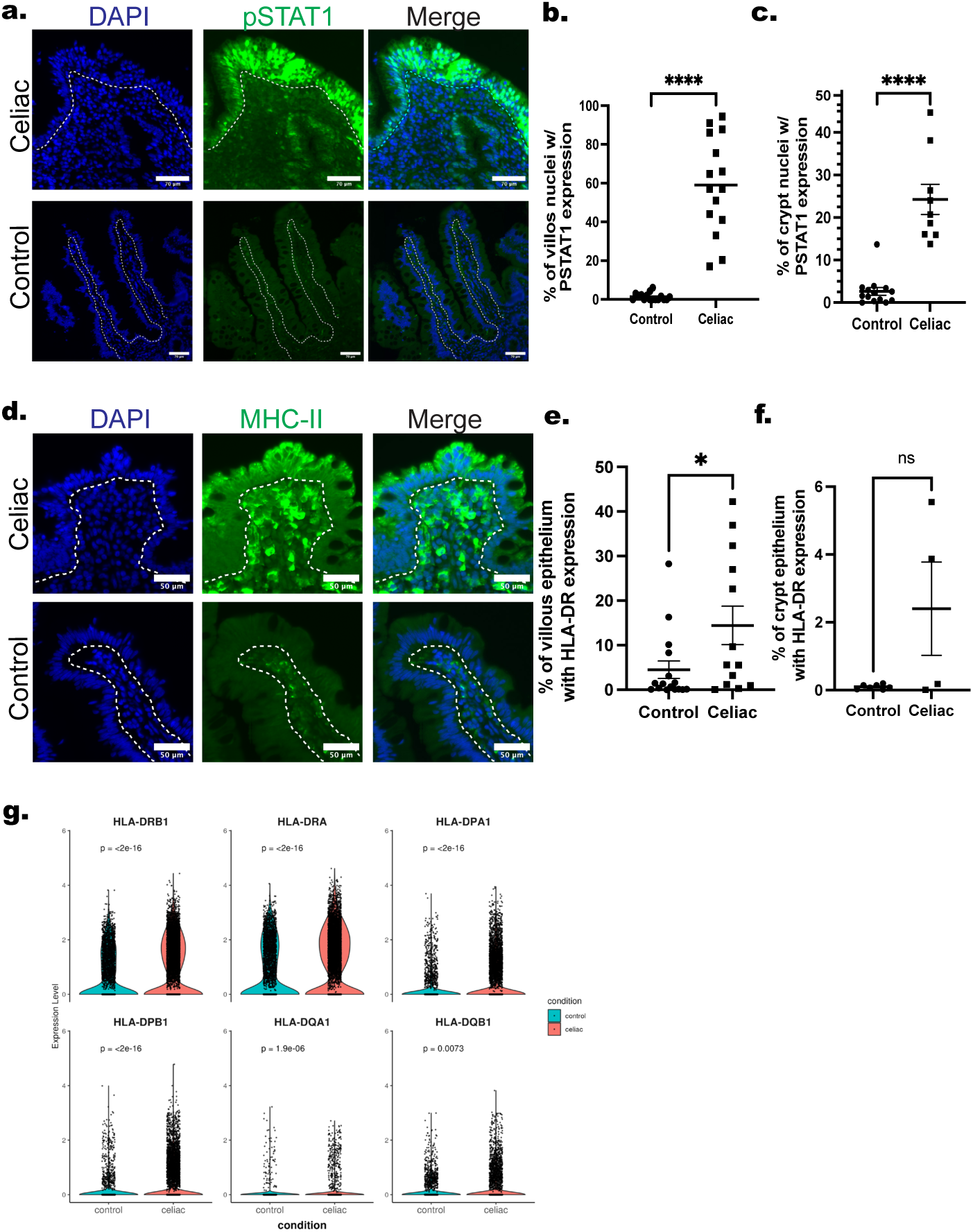
Evidence that villous enterocytes in CeD but not controls display IFNγ-mediated signaling. **(a).** CeD and control biopsies were labeled with pSTAT1 antibody. Images show pSTAT1 (green) expression in control and CeD biopsies **(b, c).** The percentage of villous or crypt nuclei positive for pSTAT1 was calculated (N = 15) (**d)**. Representative images of HLA-DR (green) expression in control and CeD biopsies **(e, f).** Quantification of HLA-DR expression on villous and crypt enterocytes in CeD and control biopsies determined by calculating the percent of HLA-DR expression normalized to epithelial area (N = 11) **(g).** Expression values of HLA Class II genes in enterocytes from control vs celiac groups in scRNA-seq data, displayed as violin plots. Significance determined by t-test. \**P* ≤ 0.05; \*\**P* ≤ 0.01; \*\*\*\**P* ≤ 0.001.

Although nuclear pSTAT1 is the hallmark of IFNγ signaling, it is not absolutely specific for IFNγ activation. IFNγ is the main human cytokine known to induce class II HLA molecules, such as HLA-DR, on epithelial cell types such as enterocytes (46). Strikingly, HLA-DR is also increased on villous enterocytes in CeD (Fig. 5d, e). This response is heterogenous, with some biopsies showing essentially all of the enterocytes expressing HLA-DR protein whereas others overlap with the controls. HLA-DR expression was also slightly elevated in the crypt (Fig. 5f). These findings are also consistent with scRNA-seq analysis of enterocytes in CeD that revealed an increase in class II HLA transcripts in villous epithelial cells from celiac biopsies (Fig. 5g). The transcript levels of HLA-DP and HLA-DQ were markedly lower than for HLA-DR. IF staining for HLA-DQ protein on enterocytes was negative (data not shown). It is unclear what role HLA-DR expression on villous enterocytes may play in CeD pathogenesis, but it supports the hypothesis that villous enterocytes are a key target of IFNγ produced by intraepithelial CTLs in CeD.

### Evidence supporting IFNγ in the recruitment of CTLs into the villous epithelium

Next, we explored two specific possible pathogenic consequences of IFNγ actions on enterocytes of relevance to CeD: CTL recruitment and CTL triggering. First, to explain the mechanism underlying the increased ratio of CTL to enterocytes within the epithelium, we examined changes in enterocyte chemokines and corresponding chemokine receptors on CTLs. To do so, we used RNAScope to probe for mRNA encoding chemokines known to recruit CTL in other settings, namely CCL3, CCL4, CCL5, CXCL9, CXC10, CXCL11, and CCL25 (47). The levels of mRNA species CCL3, CCL4, and CCL5 were significantly upregulated in the villous epithelial layer of CeD biopsies compared to controls (Table 1a, Fig. 6a-c). In our scRNA-seq experiments, CCL5 is only minimally expressed by CeD enterocytes, but it is expressed by the activated T cell populations in both CeD and controls. RNAScope also suggested that T cells may be the major source of CCL5 whereas CCL3 and CCL4 primarily localize to enterocytes of the villus so that both cell types may contribute to CTL recruitment. CXCL10 and CXCL11, were also upregulated in CeD biopsies (Fig. 6d, e). Interestingly, CXCL9, is expressed at much lower levels and is only slightly increased in CeD samples. Enterocytes also expressed mRNA for CCL25, thought to be specific for homing of T cells to the small intestine, but the level was only slightly increased in CeD vs. control biopsies (Fig. 6f). Qualitatively similar findings for these chemokines are seen by scRNA-seq analysis of isolated APOA4+ expressing enterocytes, but there appear to be quantitative differences between these techniques that may reflect specimen variation or differing sensitivity of specific transcripts to sequencing versus hybridization.

**Figure 6.**
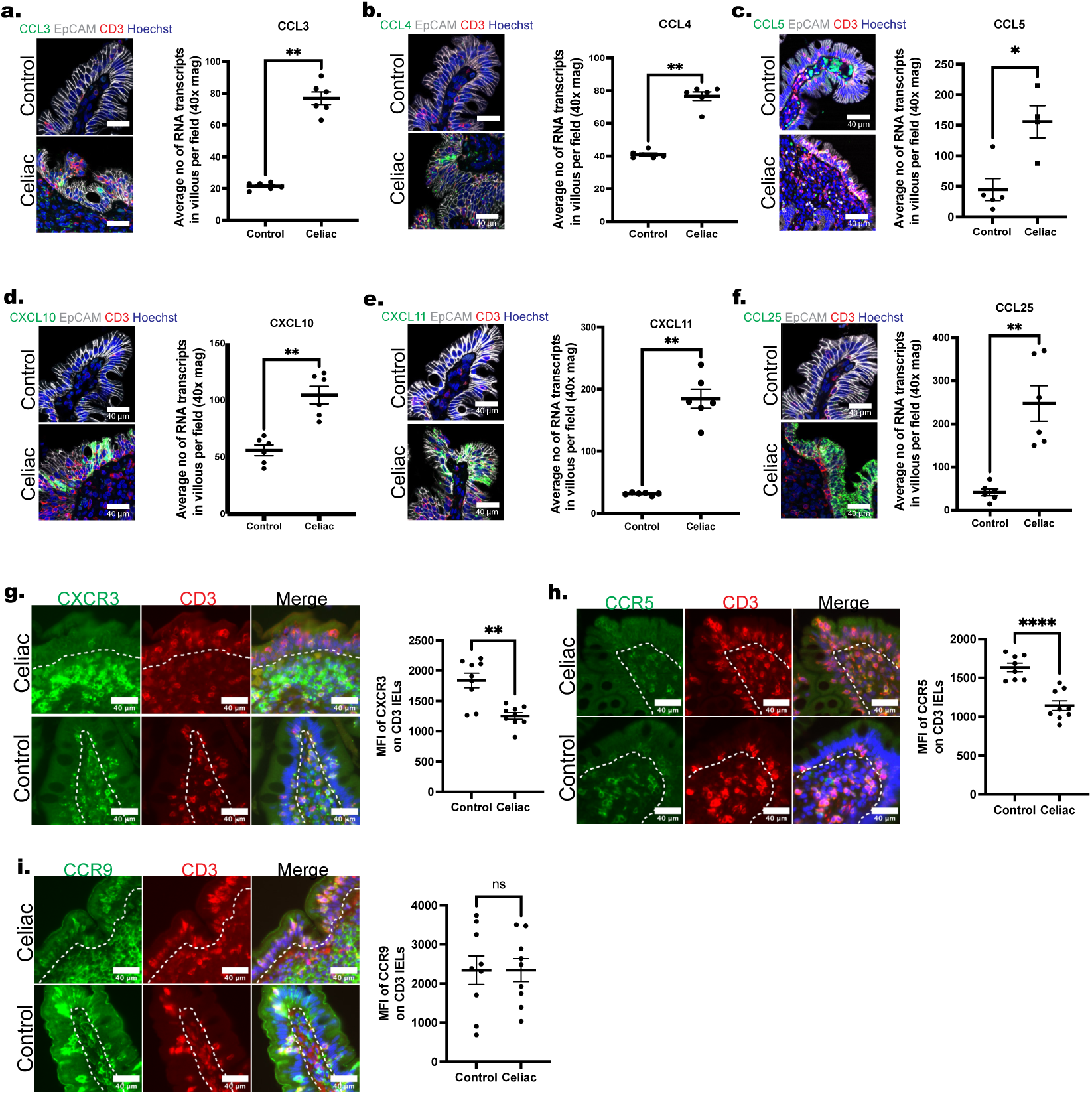
Chemokine receptor protein and chemokine mRNA expression in CeD and control biopsies. Control and celiac biopsies were stained with CD3 (red) and chemokine receptors (green) CXCR3, CCR5, and CCR9. Expression of each chemokine protein receptor was determined by calculating the mean fluorescence intensity (MFI) of **(a).** CXCR3 (N = 9), **(b).** CCR5 (N = 8), and **(c).** CCR9 (N = 9) among all CD3 cells in the villous. **(d-i)** RNAScope was used to determine expression of chemokine transcripts CCL3 (N = 6), CCL4 (N = 6), CCL5 (N = 5), CXCL10 (N = 6), CXCL11 (N = 6), and CCL25 (N=6) by calculating the average number of transcripts in the villous epithelium of CeD and control biopsies per field at 40X (N = 6). IF is used for cellular segmentation of transcript expression of chemokines (green) with epithelium labeled using EPCAM (white) and T cells with CD3 (red) in the villous of biopsies. Significance determined by t-test. \**P* ≤ 0.05; \*\**P* ≤ 0.01; \*\*\*\**P* ≤ 0.001.

By IF we found that T cells within the surface epithelium of CeD biopsies expressed CXCR3 and CCR5. (Fig. 6g, h). Chemokine receptor expression was similar in TCR αβ and γδ cells within the villous endothelium (Supplementary Fig. 4). Intraepithelial CTLs also expressed CCR9, the receptor for CCL25 (Fig. 6i).

To link changes in chemokine expression to the actions of IFNγ, we used organ cultures of duodenal mucosa from resected small bowel samples. The addition of IFNγ to organ culture led to upregulation of enterocyte nuclear pSTAT1 similar to that seen in CeD biopsies (Supplementary Fig. 6a). RNAScope analyses of the organ cultures also revealed a similar pattern to biopsies with increases in chemokine transcripts CCL3, CCL4, CXCL10, CXCL11, and CCL25 in enterocytes using antibody to EpCAM for more precise segmentation. CXCL9, however, was only slightly increased (Supplementary Table 4, Supplementary Fig. 5). However, in contrast to CeD biospies, crypt enterocytes showed similar changes as villous enterocytes when exposed to exogenous IFNγ. These data are consistent with the localization of IFNγ production to the villous layer in CeD biopsies. Collectively, our findings suggest an interpretation in which locally produced IFNγ, i.e. from intraepithelial CTL within the residual villi, induces villous enterocytes to produce chemokines that recruit CTL and that intraepithelial CTL within the villi express receptors capable of responding to these chemokines, setting off a feed forward loop when the CTLs are activated.

### Evidence for IFN in CTL recognition of enterocytes

In a final series of experiments we assessed effects of IFNγ on potential target molecules expressed on enterocytes that could activate and trigger killing by CTL. Conventional CTL target peptides bound to class I HLA-A, HLA-B or (less often) HLA-C molecules. Polymorphisms in the region of these proteins that bind 9-11 amino acid peptides determine which peptides can bind and typically only a few structurally related peptides can bind to each of the six class I inherited alleles (i.e. one HLA-A, B or C allele on each chromosome 6) (48). Although HLA-E has only limited polymorphism (only two common alleles that bind similar peptides), it too can present peptides to CTL, albeit from a very limited number of pathogens (49). NKG2C also recognizes HLA-E as a target in a manner largely independent of peptide, although some peptides when bound to HLA-E may prevent NKG2C binding (50). Thus, we first evaluated for changes of expression of these target molecules and their potential receptors on CTL in CeD biopsies vs. controls. mRNAs encoding the heavy chains of HLA-B, C and E are upregulated by IFNγ as is mRNA encoding the shared common light chain, b_2_-microglobulin. Our scRNA-seq data confirmed that transcripts encoding these proteins are elevated in CeD. HLA-A transcription, which is less responsive to IFNγ (51), was not significantly upregulated. We then used IF to examine changes in expression levels of HLA-B, HLA-E and b_2_-microglobulin protein. (We do not have a pan-HLA-A or -C reagent.) HLA-B and b_2_-microglobulin protein levels are strongly expressed in control biopsies. Although some CeD specimens showed a further increase in HLA-B expression, the overall increase in protein levels detected by staining did not reach statistical significance (Fig. 7b, c). b_2_-microglobulin expression on surface epithelium was also not significantly different from controls (Fig. 7d, e). In contrast, as previously described by others (35), HLA-E protein expression was significantly elevated in both the villous and crypt epithelium of CeD biopsies (Fig. 7f, g). We then turned to evaluate expression of potential receptors on T cells. While almost all intraepithelial CD3 T cells coexpress either αβ or γδ TCRs, fewer than 5% of intraepithelial CD3 T cells express NKG2C (Fig. 7h, i) and virtually no intraepithelial T cells in our CeD biopsies express the NKG2C-associated signaling protein DAP12 although staining is readily detectable in various L cell populations in the same biopsies (Fig. 7h, i). The latter findings are consistent with our scRNA-seq dataset which actually show a reduced number of *TYROBP* (DAP12) expressing T cells in celiac vs. control groups (Fig. 7j).

**Figure 7.**
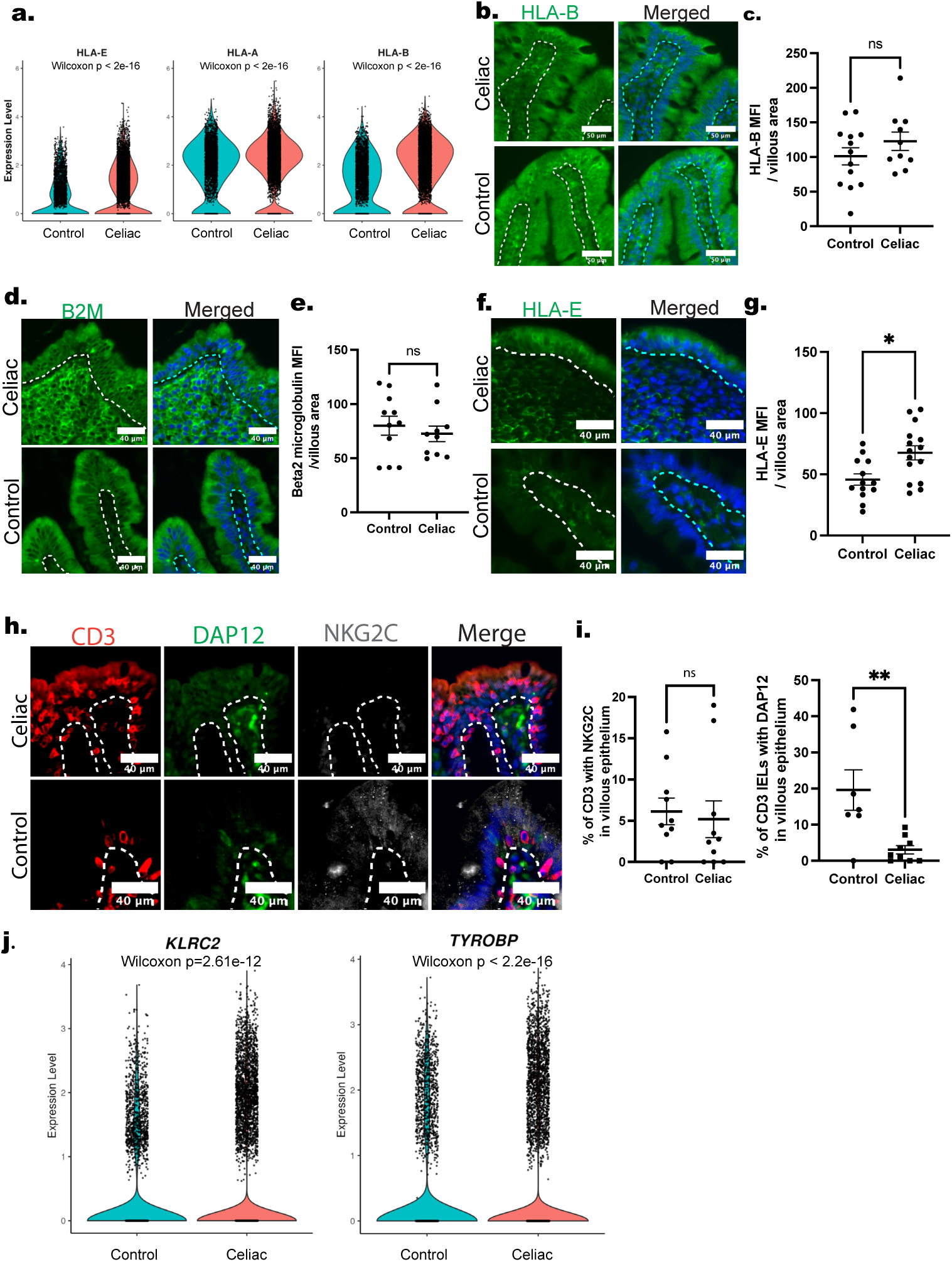
Assessment of HLA expression on villous enterocytes in CeD and controls. **(a).** Violin plots showing expression levels of HLA-E, HLA-A, and HLA-B by scRNA-seq in enterocytes. **(b-g)** Representative images and quantification of HLA-B (N=12), β2-microglobulin (N=11), and HLA-E (N=12) protein expression in the villous epithelium of CeD and control biopsies. **(h, i)**, NKG2C (N= 8) and DAP12 (N = 8) protein expression by IF in CeD and control biopsies. **(j).** Violin plots displaying expression values of KLRC2 and TYROBP (DAP12) in T cells from scRNA-seq data. Significance determined by t-test. \**P* ≤ 0.05; \*\**P* ≤ 0.01; \*\*\*\**P* ≤ 0.001.

### Activated CTL use their TCRs to engage enterocytes through HLA-B and HLA-E

Increased MHC class I molecules namely HLA-B and HLA-E on enterocytes, but lack of evidence (or weak evidence) for NKG2C signaling led us to revisit CTL-mediated interactions for killing. To initially assess the frequency to which T cells use their TCR vs activating NKG2C receptor to recognize MHC-I ligands HLA-B and HLA-E for killing, we revisited our scRNA-seq dataset and employed NICHES to analyze cell-cell interactions (52). NICHES uses ligand-receptor signal markers to identify and predict different signaling patterns within cell groups and across various experimental conditions. NICHES can operate with any list of ligand-receptor signaling mechanisms and provides a flexible analysis of cell-cell signaling at single-cell resolution. Using NICHES, we first identified top potential interacting partners between enterocytes and activated CD8 T cells in control and celiac biopsies (Supplementary Fig. 6a). These included multiple interactions that centered around HLA-B, HLA-E, and IFNG. In support of our prior findings, we found increased ligand-receptor interactions between IFNG and IFNGR1 in CeD samples (Supplementary Fig. 6b). We then focused on what molecules HLA-E could potentially interact. Interestingly, predicted interactions between enterocytes and activated CD8+ T cells were enriched in TCR-related genes such as CD8A, CD8B, and CD3D (Supplementary Fig. 6c, d). The potential interaction between HLA-E and the NK cell receptor KLRC2 (encoding NKG2C) was not significant (Supplementary Fig. 6d).

To directly assess if TCRs or NKG2C on CTLs have engaged class I HLA molecules on enterocytes, we employed PLA to detect protein-protein interactions. Because this assay only detects targets interacting that are less <40 nm apart, it can sensitively capture and quantify binding interactions between receptors and ligands expressed by interacting cells (53). As a control, we also assessed the interaction between TCRs and HLA-DR within the epithelium which, as expected, was rarely observed (Fig. 8a-d). TCR αβ and γδ cells both showed increased contacts with enterocyte MHC-I molecules in CeD biopsies compared to minimal interactions with HLA molecules in controls. The relative frequencies of such interactions, however, varied considerably (Fig. 8e-n, Table 1b). Overall, ∼33% of CD3 T cells appeared to use their TCR to interact with HLA-B or HLA-E compared to <5% in controls, and the number of TCR contacts with class I HLA molecules could be larger if we had been able to perform PLA with HLA-A and HLA-C. Importantly, T cells that used their TCRs to engage enterocyte class I MHC molecules frequently expressed Nur77, indicative of TCR signaling (54). NKG2C interactions with HLA-E were relatively rare, involving only about 3% of T cells, which is much lower than the frequency of TCR engagement with class I HLA molecules and consistent with the relatively low expression of this molecule on the intraepithelial T cell population. Thus both our computational and experimental approaches suggest that the major event triggering killing of enterocytes is likely to be mediated by TCR recognition of peptides bound conventional class I HLA molecules and, unexpectedly, to HLA-E.

**Figure 8.**
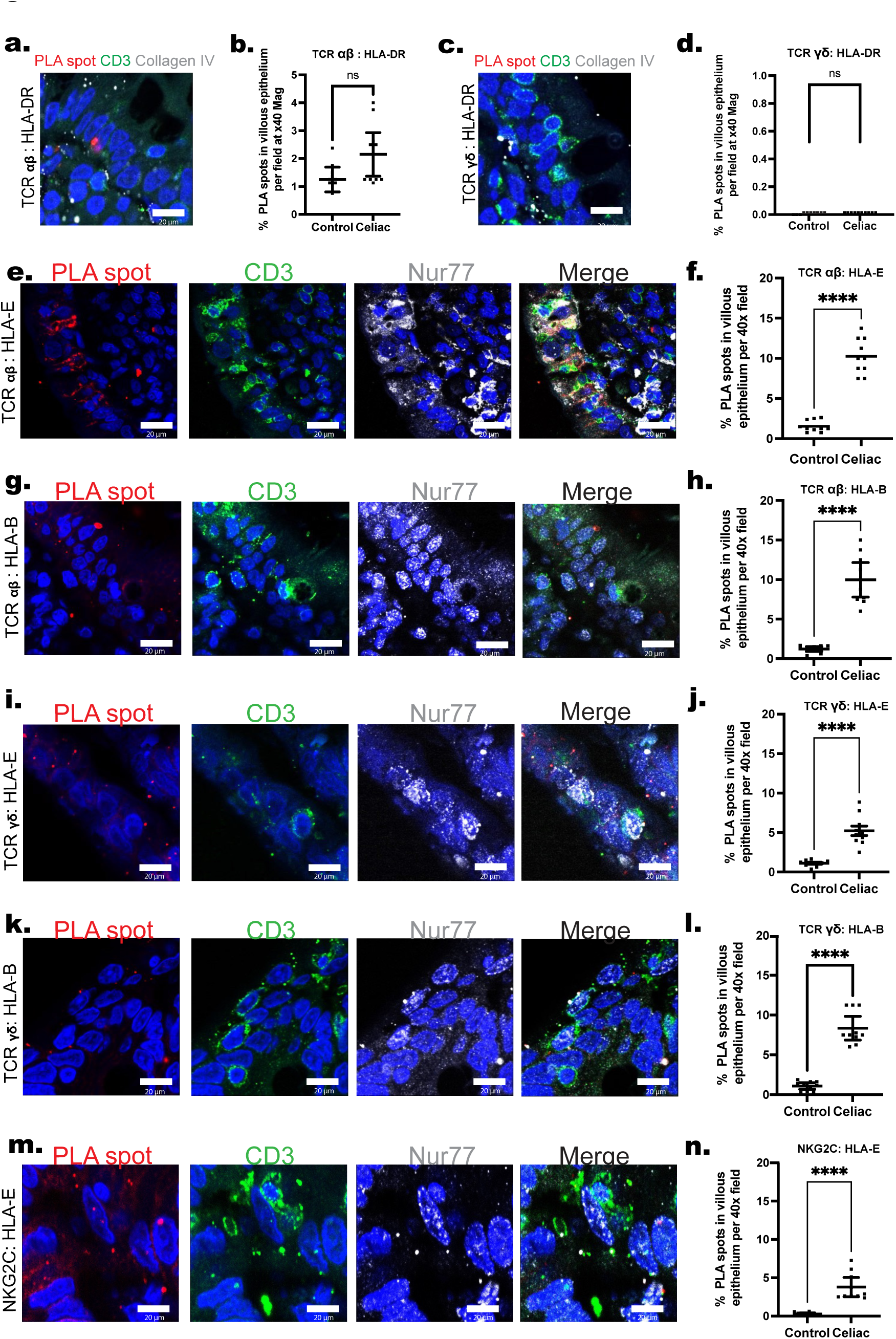
Interactions between TCRs and NKG2C on intraepithelial CTL with enterocyte class I HLA molecules. Proximity ligation assays were used to estimate the frequency of interactions between receptors TCRαβ, TCRγδ, and NKG2C to different MHC-I ligands in the villous epithelium. Biopsies were co-labeled with CD3 (green), Collagen IV (white), and counterstained with Hoechst (blue) to identify intraepithelial T cells. PLA spots (red) on images show binding between receptors and ligands. Data represent the percentage of CD3 cells with PLA spots per multiple 40x fields. (**a-d).** Binding of TCR αβ or γδ with HLA-DR was assessed as a control. Graphs quantify close interactions of TCR αβ to (**e, f)** HLA-E (N = 6) or **(g, h)** HLA-B (N = 6); TCR γδ to **(i, j)** HLA-E (N = 6) or **(k, l)** HLA-B (N=6); and NKG2C to **(m,n)** HLA-E (N=6). Significance determined by t-test. \**P* ≤ 0.05; \*\**P* ≤ 0.01; \*\*\*\**P* ≤ 0.001.

## Discussion

In this study, we used fresh and fixed duodenal mucosal biopsies from CeD patients vs. controls to characterized molecular interactions between intraepithelial T cells and enterocytes by using a combination of scRNA-seq, IF/confocal microscopy, and fluorescence in situ hybridization (RNAScope). Consistent with the literature, we saw increased ratios of intraepithelial T cells to enterocytes with many of these T cells resembling CTLs expressing CD45RO and granzyme B (18). Moreover, they also express Nur77 and Ki-67 protein as well as *IFNG* transcript indicative of T cell activation. *IFNG*-expressing T cells were less frequent in the crypts and rare in the LP, confirming a prior report that intraepithelial CTLs in the villi appear to be the major source of IFNγ in the duodenal mucosa in CeD (55). The findings within the crypt epithelium need to be interpreted with caution because crypts show regional differences in their epithelial cell linings. Thus an effect of IFNγ on some epithelial populations could be relevant for crypt elongation even if net levels of the cytokine within the crypt do not increase. It is puzzling that we did not see any significant levels of transcript encoding this cytokine in the CD4 T cell population of the LP, since these cells make IFNγ as well as IL-21 when isolated and activated in vitro (11–13). These findings are in line with a current report that transcriptionally characterized epithelial-T cell interactions in 21 CeD patients(44). In tjs study, as in ours, CD8⁺ effector memory T cells appeared to be the primary producers of IFNγ, whereas only a few CD4⁺ T cells expressed the *IFNG* transcript. CD4⁺ T cells that can be stimulated to make both IL-21 and IFNγ and are best classified as Tph cells, a population initially identified in the rheumatoid arthritis synovium and later confirmed as a major population in celiac disease and other autoimmune disorders (14, 15, 56). In nasal polyps, Tph cells require a second signal, such as IL-12 to make IFNγ (57). Perhaps such a second signal is lacking within the LP of celiac patients that can be provided by myeloid cells in peripheral blood. Alternatively, LP Tph may express IFNγ transiently, e.g., following gluten ingestion.

Due to transcriptional and protein changes strongly reflecting IFNγ signaling in villous enterocytes, we focused on the role of this cytokine (58). Specifically, we observed increased levels of nuclear pSTAT1, the principal proximal transcription factor activated by IFNγ signaling, in CeD tissue that was not observed in duodenal enterocytes in normal biopsies. The cell-cell interaction analysis indicated that activated CD8 T cells likely provided IFNG. IFNγ has been previously reported to be upregulated in CeD and a recent study bioinformatically identified IFNG and IFNGR1 as a significantly enriched ligand receptor pair in CeD (22, 44, 59). Our data specifically highlights its strong and specific effects on villous enterocytes. The plus/minus difference between villous nuclear pSTAT1 expression raises the possibility of developing this finding as a biomarker to monitor disease activity, although this must be validated in larger studies.

One potentially pathogenic role of IFNγ on villous enterocytes involves recruitment of CTLs selectively into the villous epithelium. It has been reported that upon gluten challenge in CeD, there is a rapid increase in the level of gluten-reactive CD4 T cells, CD8 αβ T cells, and γδ T cells in the circulation and a concomitant rise of T cells with overlapping TCRs in the duodenal wall (23, 27). It is likely that chemokines drive selective recruitment of circulating T cells to the duodenum, and specifically for CD8 T cells, to the epithelium. As potential evidence of IFNγ involvement as a trigger of CTL recruitment, we found elevated transcript levels of chemokines that bind CCR5 and CXCR3 produced by villous epithelium in CeD and that these same chemokines are induced by the actions of IFNγ in organ culture. Both CCR5 and CXCR3 are expressed on the intraepithelial CTL. The distribution of mRNAs coding for CXCL10 and CXCL11, which were more increased in the villous enterocytes compared to crypt enterocytes, suggest it could provide a specific recruitment signal to the villous epithelial lining rather than the epithelial lining of the crypts. We do not know if crypt enterocytes are less responsive to IFNγ than are villous enterocytes or whether the lesser ratio of intraepithelial T cells to enterocytes in the crypts vs. the villi means the IFNγ concentration is likely lower in the crypts than in the villi. A third ligand for CCR5, namely CCL5, is also upregulated in CeD, but it is produced more prominently by activated intraepithelial T cells than by enterocytes and there are fewer T cells in the crypts than the villi. Moreover, our organ culture experiments, performed using normal duodenal mucosa, suggest thet crypt enterocytes do respond to IFNγ. At the same time we speculate that CCL25 acts on circulating T cells bearing CCR9 for the initial phase of recruitment from blood into the LP (60).

The second major implication of our study is an assessment of how IFNγ may contribute to villus enterocyte loss. Assuming that CTLs are killing villous enterocytes based on recognition of a signal expressed by enterocytes in CeD, we used both NICHES and PLAs to evaluate two different theories of T cell mediated killing in CeD: 1). TCR-mediated killing that is antigen driven with recognition of antigen (likely gluten peptides) on classical MHC-1 molecules and 2). NKG2C receptor mediated cytotoxicity that is non-antigen specific. Transcription of HLA-B and HLA-E heavy chains as well as b_2_-microglobulin are all known to be upregulated by IFNγ (61, 62). However, only HLA-E protein is strongly induced above control levels in CeD. The discrepancy between mRNA and protein may be explained by the already significant levels of HLA-B in the controls. The increase of HLA-E above controls corroborates other studies that have similarly found elevated HLA-E levels on villous enterocytes in CeD (35, 44). The increase in HLA-E expression has been cited to support the hypothesis that HLA-E recognition by NKG2C, a natural killer (NK) receptor that can be induced on highly activated CTLs, bypasses the need for a TCR signal (35). However, our tissue analysis detected an average of fewer than 5% of intraepithelial T cells expressing NKG2C. Similarly, a recent study by FitzPatrick et al., observed only a small subset of CD8+ T cells expressing NKG2C (around 20%). Intereastingly, NKG2C was more abundant in natural CD8+ IELs, which only make up a very small subset of CD8+ T cells in CeD as they are replaced with CTL. Moreover, we did not detect expression of DAP12 in intraepithelial CTL, although it is readily detected within cells of the LP. Another report utilizing scRNA-seq also failed to see DAP12 expression in CeD intraepithelial T cells (22). While our own scRNA-seq experiments did show some DAP12 expression, the levels did not differ significantly between CeD and control biopsies. We are uncertain whether the lesser expression of NKG2C and DAP12 we see arises from a difference in sensitivity of detection by staining of intact biopsies or from our ability to exclude LP T cells from our spatial but not scRNAseq analyses.

Our application of NICHES to our scRNAseq analysis supported the hypothesis that both HLA-B and HLA-E were interacting with TCRs of the intraepithelial CTL in CeD biopsies. Surprisingly, most of the TCR γδ CTL in our scRNAseq analysis also expressed CD8 and could be class I HLA restricted. In our final set of experiments, we used PLA to directly identify the interactions of intraepithelial CTL receptors with enterocyte class I HLA molecules. The most abundant interactions involved αβ TCRs with HLA-E followed by interactions with HLA-B. γδ TCRs also interacted with HLA-E and HLA-B. We did not have reagents to test interactions with HLA-A or -C, but we expect that these class I molecules could also be relevant. As a negative control, we tested for interactions between CTL TCRs with HLA-DR, an I-induced class II MHC molecule associated with peptide presentation to CD4 T cells, and found no significant contacts. Additionally, we found no significant level of TCR interactions with class I HLA molecules in the control biopsies. We did find NKG2C interactions with HLA-E, but consistent with its infrequent expression on T cells, NKG2C interactions were relatively uncommon compared to TCR interactions. Our interpretation that intraepithelial CTLs are responding to antigens presented by class I HLA molecules is consistent with some recent data indicating that CTL, like CD4 cells, can respond to some gluten-derived peptides, but such peptides need not be deamidated (29–31). TCR involvement in CeD is supported by recent single-cell and bulk duodenal CD8+ T cell TCR analyses, which demonstrate clonal enrichment of specific TCRs in the CeD duodenum—a feature not observed in non-CeD inflammatory gastrointestinal conditions (44), suggesting involvement of antigen. However, these studies did not determine the antigen specificity of these TCRs.

Our data point away from a primary role for NKG2C interactions with HLA-E but do not negate a role for other NK receptors on T cells, such as NKG2D, interacting with enterocyte ligands (63). However, in humans, unlike mice, NKG2D cannot signal through DAP12 and primarily functions to increase the responsiveness to signals like TCR engagement rather than directly trigger CTL-mediated killing (64). It remains to be determined what other costimulators on enterocytes could be contributing to CTL killing.

We are uncertain how enterocytes are able to load gliadin peptides, which are derived from an exogenous protein, on to HLA class I proteins. Such peptides are normally presented by class II HLA molecules. Dendritic cells are known to be able to load such peptides on the conventional class I molecules, a process called cross-presentation (65). It is not known if enterocytes may also cross-present. Alternatively, perhaps extracellular peptides simply displace self peptides on surface expressed class I HLA proteins. The argument against this possibility is that class I HLA molecules are selective for peptides of a specific length (9-11 amino acids) and cannot bind larger or smaller fragments and the intracellular processing machinery functions to generate such fragments whereas random proteolysis does not.

The increased interactions of TCR molecules to not only classical HLA class I molecules but also HLA-E in CeD was unexpected and warrants further discussion. HLA-E proteins are intrinsically unstable in the absence of bound peptide and very few peptides are known to bind. Unstable class I molecules, including HLA-E, are typically retained and degraded in the endoplasmic reticulum. Translocation of HLA-E to the cell surface requires binding of self-peptides most often derived from the signal sequences of other HLA class I molecules, especially HLA-A2 (66), but such peptides are limited in quantity, typically resulting in very low cell surface expression of HLA-E at baseline (38). Furthermore, those HLA-E molecules that do reach the cell surface are rapidly endocytosed, a process selective for HLA-E vs. other class I proteins, further limiting surface expression. Despite its limited polymorphism, the HLA-E proteins have been shown to present peptides from certain infectious agents, including *Mycobacterium tuberculosis*, Human Immunodeficiency Virus and influenza to TCRs (50, 67). These microbial peptides are delivered to and bind HLA-E molecules only after HLA-E proteins have reached the surface and have then been internalized into an endocytic compartment. Binding of these exogenous peptides leads to stable re-expression of HLA-E on the cell surface and presentation to CD8+ CTL. In other words, there must be a source of such peptides in order to observe increased HLA-E expression. Importantly, some gliadin-derived peptides have been shown to similarly stabilize and increase HLA-E expression in dendritic cells and may be important in the upregulation of HLA-E on enterocytes seen in CeD (68). Rather than classify HLA-E as a non-conventional class I HLA molecule or as a “stress-induced protein,” it may be more accurate to consider HLA-E as a “specialist” HLA class I molecule similar to the BF1 locus in chickens (69). If this interpretation is correct, then a key element of CeD pathogenesis is the ability of certain gliadin peptides to mimic serious pathogen-derived peptides and increase HLA-E expression.

There are some limitations of this study. First, it is possible that CeD is not a single entity but multiple entities that have different endotypes. Our sample size is not large enough to detect this, but our findings so far do not suggest significant differences among the patients studied that correlate with sex or age. Our PLAs do not establish that the molecular interactions we detect are gliadin peptide-dependent or result in TCR signaling. However, we do see Nur77 expression in the same T cells, consistent with activation by TCR engagement (70). Despite these limitations, we believe that this study has provided important evidence to support a testable role of gluten peptide antigen recognition as the primary mechanism by which intraepithelial CTLs are recruited and target villous epithelial architecture and function. Specifically, the central roles of IFNγ and HLA-E suggested by these studies, which defines potentially relevant actions of this cytokine in CeD, provides a path for both new biomarker and therapeutic development.

## Materials and Methods

### Sex as a biological variable

Our study examined tissue samples from male and female patients, and similar findings were found in both sexes.

### Tissue processing and single cell droplet encapsulation

Duodenal mucosal biopsies for research were obtained with informed consent from 3 patients with CeD with mild villous blunting (Marsh grade 3A) or severe villous blunting (Marsh grade 3C) and 3 patients with normal mucosa undergoing diagnostic endoscopy. All six samples were deidentified prior to transfer to the laboratory under a protocol approved by the Yale Institutional Review Board. Research biopsies were immediately immersed in RPMI containing 1 mM DTT, 20 mM EDTA and 2% FBS at 37° C for 15 min to isolate epithelial cells and dissociated in Liberase TL (200 μg/ml of Liberase TL) containing solution in RPMI for 20 min at 37°C. Digested cell suspensions were passed through a 100-μm cell strainer at least three times, washed with complete media, and combined with the epithelial fraction before processing for droplet-based single-cell RNA-seq (scRNA-seq). To ensure 10,000 cells were encapsulated into droplets using 10x Chromium GEM technology single cells were suspended at 1,500 cells/ microliter concentration. Libraries were prepared in house using the Chromium Next GEM Single Cell 3′ Reagent Kits v.3.1 (10x Genomics). scRNA-seq libraries were sequenced using the Nova-Seq system.

### Immunofluorescence analysis of duodenal mucosa

All archived human tissue samples used in these studies were de-identified and declared as not human subjects research by the Yale Institutional Review Board. Formalin fixed paraffin embedded (FFPE) duodenal mucosal diagnostic biopsies had been obtained from CeD positive cases and/or from patients in whom CeD and other pathological changes were ruled out (normal controls), all accessed through the Yale Department of Pathology. Because the degree of morphological severity can vary at different sites within the duodenum of the same CeD patient, common diagnostic practice is to take multiple duodenal mucosal biopsies during a single endoscopy. The Marsh-Oberhuber Score is used as an index to classify different stages of tissue injury in celiac disease with score 3C being the most severe, showing complete villous blunting, crypt hyperplasia, and increased ratios of intraepithelial T cells to enterocytes (71). For analysis of FFPE CeD biopsies in this study, we selected biopsy specimens with a Marsh-Oberhuber Score of 3B-3C. The sex and age of the patient and control samples analyzed are summarized in Supplementary Table 1. For each parameter assessed, archival FFPE biopsies from CeD patients and normal controls (i.e. no pathological changes identified) were screened by a pathologist (MER) to identify relevant tissue fragments and then re-embedded as individual samples or as multiple (5-6 mm) CeD and control samples from different patients displayed on a single slide as a tissue microarray (TMA) (Supplementary Table 2). For analysis, the data from multiple samples are pooled for each parameter examined and the numbers of different biopsies analyzed for that parameter are noted in the legend of each figure. One TMA was also created using paired biopsies from four CeD patients who had tissue sampled at the time of diagnosis and later after symptom abatement on a gluten-free diet.

For IF staining, tissues were deparaffinized with xylene and hydrated with graded washes in ethanol. Antigen retrieval was then performed using citrate-based antigen unmasking solution (Vector Laboratories, H-3300) with samples either heated to 95°C in solution on a hot plate for 30 minutes or using a pressure cooker for 5 minutes. Sections were incubated with blocking buffer (5% donkey or goat serum in PBS, 1% BSA-antibody dilution buffer) for 1hr at room temperature (RT) to block nonspecific binding of antibodies to tissue. Primary antibodies were then added overnight at 4°C (Supplementary Table 3), then incubated with appropriate AlexaFluor-conjugated secondary antibodies in blocking buffer (1:200) for 1hr at RT. All antibodies used were validated by the vendor and by using tonsil, spleen, and/or Crohn’s disease tissue to confirm reactivity and specificity. Secondary only and isotype controls were used to distinguish signal from background fluorescence.

### Cyclic Immunofluorescence (CycIF) of FFPE tissue

CycIF involves repeated staining and imaging of the same tissue section by quenching the fluorescent signal and adding directly conjugated antibody (72). This was performed manually in the present study. After tissue staining, coverslips are placed on slides using 10% glycerol in PBS. Once images have been acquired, coverslips are removed by placing slides in PBS for 30 min, followed by bleaching the sections with 3% H_2_O_2_ and 20mM NaOH in PBS for 1 hr while exposed to light to quench the fluorescence. After thorough washes in PBS, specimens are then restained with conjugated antibodies overnight. Images are recorded at each step and then brought into spatial alignment for colocalization of staining patterns using ImageJ and fluorescence intensity quantified by QuPath (72).

### Combined IF and In Situ Proximity Ligation Assay (PLA)

A NaveniFlex^TM^ Tissue MR (Red/Atto647) PLA kit from Navinci Diagnostics (Uppsala, Sweden) was used following the manufacturer’s instructions. Briefly, following dewaxing and hydration, sections were incubated with a combination of two primary antibodies, either against a 1). TCR molecule (αβ or γδ) and MHC ligand (HLA-E, HLA-B or HLA-DR) or 2). NKG2C and HLA-E overnight at 4°C. Navenibody M1 and R2 probes were then added for 60 min at 37°C, followed by two enzymatic reactions to create a rolling circle amplification. At the post-block stage, primary antibodies to CD3 and Collagen IV (to mark basement membrane of the villous epithelium) were added for 30 minutes, followed by incubation with the detection reagent (Atto647) and fluorescence-conjugated secondary antibodies for 60 min at 37°C. After mounting with Prolong Gold Antifade mountant (P36930, Thermo Fisher) and counterstaining the nuclei with DAPI, PLA red fluorescence spots in villous epithelium, indicative for protein-protein interaction, were evaluated using the confocal microscope. The average percentage of PLA spots in villous epithelium in ten fields of view at 40x magnification was counted in ImageJ and quantified using GraphPad Prism. For negative controls, the primary antibody was replaced with isotype-specific serum and IF staining with the antibodies of the PLA pair were included in parallel with the PLA as a control to demonstrate effective binding by the primary antibodies in each experiment.

### Duodenal Mucosal Organ culture

Normal duodenal mucosa was provided by the Yale Department of Pathology from excised, de-identified and discarded surgical intestinal tissue from patients undergoing Whipple procedures at Yale New Haven Hospital. Tissue was cut into small fragments and placed on a foam pad for adhesion. Samples were then cultured at 95% O_2_ in a 24-well plate submerged with DMEM supplemented with 10% fetal bovine serum, 2 mM L-glutamine, and penicillin/ streptomycin (73). Samples were either left unstimulated or were activated by adding 500ng/mL of recombinant human IFNγ (PeproTech 300-02) for 24 hours, followed by fixation in 10% NBF for 24 hours and paraffin embedding.

## Author Contributions

JEJ performed the IF assays. KA performed single cell RNA-sequencing analysis. RSA performed most the RNAscope and PLA assays with the rest being done by TZ, SAT, and SJL. F.Z. perfomed scRNA-seq experiments with ES and pre-processed scRNA-seq data. WX, LR, and MER provided the tissue specimens for all the experiments. JEJ, JP, RAF, and AJM helped conceieve the experiments. JEJ, JP, ES, and KA wrote the manuscript. First authorship order was determined based on the fact that both authors contributed equally but more biological assays are included in the paper.

## Acknowledgements

Supported by a Yale School of Medicine Teams Award and grants from the Colton Center for Autoimmunity at Yale to MRE and JSP, and to ES and RAF. RAF is also supported by Howard Hughes Medical Institute and ES is also supported by a grant from the Celiac Disease Foundation. JEJ is supported by the MSTP T32GM136651 training grant. RSA-L is supported by National Institute for Health and Care Research Cambridge Biomedical Research Centre at the Cambridge University Hospitals National Health Service Foundation Trust.

## References

1. I. Parzanese et al., Celiac disease: From pathophysiology to treatment. World J Gastrointest Pathophysiol 8, 27–38 (2017).

2. N. Patel, M. E. Robert, Frontiers in Celiac Disease: Where Autoimmunity and Environment Meet. Am J Surg Pathol 46, e43–e54 (2022).

3. A. Rubio-Tapia et al., American College of Gastroenterology Guidelines Update: Diagnosis and Management of Celiac Disease. Am J Gastroenterol 118, 59–76 (2023).

4. B. Lebwohl, A. Rubio-Tapia, Epidemiology, Presentation, and Diagnosis of Celiac Disease. Gastroenterology 160, 63–75 (2021).

5. S. Husby, J. A. Murray, D. A. Katzka, AGA Clinical Practice Update on Diagnosis and Monitoring of Celiac Disease—Changing Utility of Serology and Histologic Measures: Expert Review. Gastroenterology 156, 885–889 (2019).

6. L. M. Sollid, E. Thorsby, HLA susceptibility genes in celiac disease: genetic mapping and role in pathogenesis. Gastroenterology 105, 910–922 (1993).

7. E. Gnodi, R. Meneveri, D. Barisani, Celiac disease: From genetics to epigenetics. World J Gastroenterol 28, 449–463 (2022).

8. L. M. Sollid et al., Evidence for a primary association of celiac disease to a particular HLA-DQ α/β heterodimer. Journal of Experimental Medicine 169, 345–350 (1989).

9. O. Molberg et al., Tissue transglutaminase selectively modifies gliadin peptides that are recognized by gut-derived T cells in celiac disease. Nat Med 4, 713–717 (1998).

10. Y. Van de Wal et al., Cutting Edge: Selective Deamidation by Tissue Transglutaminase Strongly Enhances Gliadin-Specific T Cell Reactivity1. The Journal of Immunology 161, 1585–1588 (1998).

11. M. Bodd et al., HLA-DQ2-restricted gluten-reactive T cells produce IL-21 but not IL-17 or IL-22. Mucosal Immunology 3, 594–601 (2010).

12. E. M. Nilsen et al., Gluten induces an intestinal cytokine response strongly dominated by interferon gamma in patients with celiac disease. Gastroenterology 115, 551–563 (1998).

13. M. A. van Leeuwen et al., Increased production of interleukin-21, but not interleukin-17A, in the small intestine characterizes pediatric celiac disease. Mucosal Immunology 6, 1202–1213 (2013).

14. K. E. Marks, D. A. Rao, T peripheral helper cells in autoimmune diseases. Immunol Rev 307, 191–202 (2022).

15. A. Christophersen et al., Distinct phenotype of CD4+ T cells driving celiac disease identified in multiple autoimmune conditions. Nature Medicine 25, 734–737 (2019).

16. V. De Re, R. Magris, R. Cannizzaro, New Insights into the Pathogenesis of Celiac Disease. Front Med (Lausanne) 4, 137 (2017).

17. M. Svensson et al., CCL25 mediates the localization of recently activated CD8alphabeta(+) lymphocytes to the small-intestinal mucosa. J Clin Invest 110, 1113–1121 (2002).

18. G. Oberhuber et al., Evidence that intestinal intraepithelial lymphocytes are activated cytotoxic T cells in celiac disease but not in giardiasis. Am J Pathol 148, 1351–1357 (1996).

19. M. Haghbin et al., The role of CXCR3 and its ligands CXCL10 and CXCL11 in the pathogenesis of celiac disease. Medicine (Baltimore) 98, e15949 (2019).

20. C. Bondar et al., Role of CXCR3/CXCL10 Axis in Immune Cell Recruitment into the Small Intestine in Celiac Disease. Plos One 9, e89068 (2014).

21. G. Pietz et al., Immunopathology of childhood celiac disease-Key role of intestinal epithelial cells. PLoS One 12, e0185025 (2017).

22. N. Atlasy et al., Single cell transcriptomic analysis of the immune cell compartment in the human small intestine and in Celiac disease. Nat Commun 13, 4920 (2022).

23. A. Christophersen et al., Pathogenic T Cells in Celiac Disease Change Phenotype on Gluten Challenge: Implications for T-Cell-Directed Therapies. Adv Sci (Weinh) 8, e2102778 (2021).

24. M. Barry, R. C. Bleackley, Cytotoxic T lymphocytes: all roads lead to death. Nat Rev Immunol 2, 401–409 (2002).

25. Z. L. Z. Hay, J. E. Slansky, Granzymes: The Molecular Executors of Immune-Mediated Cytotoxicity. Int J Mol Sci 23, 1833 (2022).

26. A. Han et al., Dietary gluten triggers concomitant activation of CD4+ and CD8+ αβ T cells and γδ T cells in celiac disease. Proc Natl Acad Sci U S A 110, 13073–13078 (2013).

27. L. F. Risnes et al., Circulating CD103(+) gammadelta and CD8(+) T cells are clonally shared with tissue-resident intraepithelial lymphocytes in celiac disease. Mucosal Immunol 14, 842–851 (2021).

28. A. Kornberg, et al., Gluten induces rapid reprogramming of natural memory αβ and γδ intraepithelial T cells to induce cytotoxicity in celiac disease. Sci Immunol 8 (2023).

29. S. Picascia et al., Gliadin-Specific CD8(+) T Cell Responses Restricted by HLA Class I A*0101 and B*0801 Molecules in Celiac Disease Patients. J Immunol 198, 1838–1845 (2017).

30. C. Gianfrani et al., Celiac Disease Association with CD8+ T Cell Responses: Identification of a Novel Gliadin-Derived HLA-A2-Restricted Epitope1. The Journal of Immunology 170, 2719–2726 (2003).

31. G. Mazzarella et al., Gliadin activates HLA class I-restricted CD8+ T cells in celiac disease intestinal mucosa and induces the enterocyte apoptosis. Gastroenterology 134, 1017–1027 (2008).

32. L. M. Eggesbø et al., Single-cell TCR sequencing of gut intraepithelial γδ T cells reveals a vast and diverse repertoire in celiac disease. Mucosal Immunology 13, 313–321 (2020).

33. L. M. Eggesbø et al., Single-cell TCR repertoire analysis reveals highly polyclonal composition of human intraepithelial CD8+ αβ T lymphocytes in untreated celiac disease. European Journal of Immunology 51, 1542–1545 (2021).

34. D. A. Yohannes et al., Deep sequencing of blood and gut T-cell receptor β-chains reveals gluten-induced immune signatures in celiac disease. Scientific Reports 7, 17977 (2017).

35. B. Meresse et al., Reprogramming of CTLs into natural killer-like cells in celiac disease. Journal of Experimental Medicine 203, 1343–1355 (2006).

36. B. Jabri et al., TCR Specificity Dictates CD94/NKG2A Expression by Human CTL. Immunity 17, 487–499 (2002).

37. V. Abadie, B. Jabri, IL-15: a central regulator of celiac disease immunopathology. Immunological Reviews 260, 221–234 (2014).

38. G. M. Gillespie, M. N. Quastel, A. J. McMichael, HLA-E: Immune Receptor Functional Mechanisms Revealed by Structural Studies. Immunological Reviews 329, e13434 (2025).

39. R. Elmentaite et al., Cells of the human intestinal tract mapped across space and time. Nature 597, 250–255 (2021).

40. J. Taavela et al., Apolipoprotein A4 Defines the Villus-Crypt Border in Duodenal Specimens for Celiac Disease Morphometry. Front Immunol 12, 713854 (2021).

41. A. Popp et al., A New Intraepithelial gammadelta T-Lymphocyte Marker for Celiac Disease Classification in Formalin-Fixed Paraffin-Embedded (FFPE) Duodenal Biopsies. Dig Dis Sci 66, 3352–3358 (2021).

42. A. Kirmizi et al., Discriminant value of IEL counts and distribution pattern through the spectrum of gluten sensitivity: a simple diagnostic approach. Virchows Archiv 473, 551–558 (2018).

43. J. F. Ludvigsson et al., Diagnosis and management of adult coeliac disease: guidelines from the British Society of Gastroenterology. Gut 63, 1210–1228 (2014).

44. M. E. B. FitzPatrick et al., Immune-epithelial-stromal networks define the cellular ecosystem of the small intestine in celiac disease. Nat Immunol 26, 947–962 (2025).

45. X. Hu, J. Li, M. Fu, X. Zhao, W. Wang, The JAK/STAT signaling pathway: from bench to clinic. Signal Transduct Target Ther 6, 402 (2021).

46. J. E. Wosen, D. Mukhopadhyay, C. Macaubas, E. D. Mellins, Epithelial MHC Class II Expression and Its Role in Antigen Presentation in the Gastrointestinal and Respiratory Tracts. Front Immunol 9, 2144 (2018).

47. P. J. Trivedi, D. H. Adams, Chemokines and Chemokine Receptors as Therapeutic Targets in Inflammatory Bowel Disease; Pitfalls and Promise. J Crohns Colitis 12, S641–S652 (2018).

48. M. Wieczorek et al., Major Histocompatibility Complex (MHC) Class I and MHC Class II Proteins: Conformational Plasticity in Antigen Presentation. Front Immunol 8, 292 (2017).

49. G. Pietra, C. Romagnani, C. Manzini, L. Moretta, M. C. Mingari, The emerging role of HLA-E-restricted CD8+ T lymphocytes in the adaptive immune response to pathogens and tumors. J Biomed Biotechnol 2010, 907092 (2010).

50. M. J. Hogan et al., Cryptic MHC-E epitope from influenza elicits a potent cytolytic T cell response. Nat Immunol 24, 1933–1946 (2023).

51. D. R. Johnson, Locus-Specific Constitutive and Cytokine-Induced HLA Class I Gene Expression The Journal of Immunology 170, 1894–1902 (2003).

52. M. S. B. Raredon et al., Comprehensive visualization of cell-cell interactions in single-cell and spatial transcriptomics with NICHES. Bioinformatics 39, btac775 (2023).

53. M. S. Alam, Proximity Ligation Assay (PLA). Curr Protoc Immunol 123(1):e58 (2018).

54. J. F. Ashouri et al., Reporters of TCR signaling identify arthritogenic T cells in murine and human autoimmune arthritis. Proceedings of the National Academy of Sciences 116, 18517–18527 (2019).

55. R. W. Olaussen et al., Interferon-gamma-secreting T cells localize to the epithelium in coeliac disease. Scand J Immunol 56, 652–664 (2002).

56. Y. Huang et al., T peripheral helper cells in autoimmune diseases: What do we know? Front Immunol 14, 1145573 (2023).

57. L. Xiao et al., Human IL-21+IFN-gamma+CD4+ T cells in nasal polyps are regulated by IL-12. Sci Rep 5, 12781 (2015).

58. V. Dotsenko et al., Transcriptomic analysis of intestine following administration of a transglutaminase 2 inhibitor to prevent gluten-induced intestinal damage in celiac disease. Nat Immunol 25, 1218–1230 (2024).

59. K. E. Lundin et al., Gliadin-specific, HLA-DQ(alpha 1*0501,beta 1*0201) restricted T cells isolated from the small intestinal mucosa of celiac disease patients. J Exp Med 178, 187–196 (1993).

60. X. Wu et al., The Roles of CCR9/CCL25 in Inflammation and Inflammation-Associated Diseases. Front Cell Dev Biol 9, 686548 (2021).

61. A. J. Collins T Fau-Korman et al., Immune interferon activates multiple class II major histocompatibility complex genes and the associated invariant chain gene in human endothelial cells and dermal fibroblasts. Proc Natl Acad Sci U S A 81, 4917–4921 (1984).

62. S. Mizuno, J. A. Trapani, B. H. Koller, B. Dupont, S. Y. Yang, Isolation and nucleotide sequence of a cDNA clone encoding a novel HLA class I gene. J Immunol 140, 4024–4030 (1988).

63. B. Meresse et al., Coordinated induction by IL15 of a TCR-independent NKG2D signaling pathway converts CTL into lymphokine-activated killer cells in celiac disease. Immunity 21, 357–366 (2004).

64. K. Prajapati, C. Perez, L. B. P. Rojas, B. Burke, J. A. Guevara-Patino, Functions of NKG2D in CD8(+) T cells: an opportunity for immunotherapy. Cell Mol Immunol 15, 470–479 (2018).

65. J. E. Grotzke, D. Sengupta, Q. Lu, P. Cresswell, The ongoing saga of the mechanism(s) of MHC class I-restricted cross-presentation. Curr Opin Immunol 46, 89–96 (2017).

66. M. H. Lampen et al., Alternative peptide repertoire of HLA-E reveals a binding motif that is strikingly similar to HLA-A2. Mol Immunol 53, 126–131 (2013).

67. Z. Hannoun et al., Identification of novel HIV-1-derived HLA-E-binding peptides. Immunol Lett 202, 65–72 (2018).

68. G. Terrazzano et al., Gliadin regulates the NK-dendritic cell cross-talk by HLA-E surface stabilization. J Immunol 179, 372–381 (2007).

69. J. Kaufman, Generalists and Specialists: A New View of How MHC Class I Molecules Fight Infectious Pathogens. Trends Immunol 39, 367–379 (2018).

70. J. F. Ashouri, A. Weiss, Endogenous Nur77 Is a Specific Indicator of Antigen Receptor Signaling in Human T and B Cells. J Immunol 198, 657–668 (2017).

71. A. Ensari, M. N. Marsh, Diagnosing celiac disease: A critical overview. Turk J Gastroenterol 30, 389–397 (2019).

72. Z. Du et al., Qualifying antibodies for image-based immune profiling and multiplexed tissue imaging. Nat Protoc 14, 2900–2930 (2019).

73. K. Tsilingiri et al., Probiotic and postbiotic activity in health and disease: comparison on a novel polarised ex-vivo organ culture model. Gut 61, 1007–1015 (2012).

