## Supplementary Tables and Figures for "T Cell Receptors for Antigen on Intraepitheial Cytolytic T Lymphocytes in Celiac Disease Engage Enterocyte HLA-E and HLA-B"

### Supporting Information Text

### SI Materials and Methods

#### Single cell RNA-sequencing analysis of human duodenal biopsies

Raw sequencing data were processed using CellRanger software (v7.1.0) and aligned to the reference genome GRCh38-2020-A. Filtered output from CellRanger was used for downstream processing using R (v4.3.3) and Seurat (v5.0.0). The analysis pipeline consisted of filtering cells based on mitochondrial gene counts less than 50%, which is consistent with other scRNA-seq studies of small intestinal tissue (1). A modified version of HTODemux pipeline was used for the a2-2024 and the celiac403 samples, which both had hashed PBMC and tissue. Cells labeled PBMCs were excluded from the analysis. Gene expression was normalized and log-transformed. Cells identified as doublets by scDblFinder (v1.16.0) were removed (2). Harmony (v1.2.0) (3) was used to compute a batch-corrected embedding of the samples and to integrate the individual patient data. After integrating the data, Celltypist (v.1.3.0) (4) and the available human gut reference (1) were used to annotate cells. These annotations were manually confirmed using known marker genes and higher level annotations were created based on intial celltypist annotations that grouping subcatergories of cells. Finally, a Wilcoxon Rank Sum Test was used to find differentially expressed genes between the celiac and control samples for each cell type individually. For the T cell subset analysis, cell types that were exclusive to a patient, defined by over 90% of cells coming from a single sample, were excluded. Single cell ligand receptor interactions were found using NICHES (5). A custom ligand receptor interaction database was created using a combination of ligand and receptor interaction pairs from the Omnipath catalog and curated intestinal and celiac specific interactions to calculate differential ligand receptor interactions between control and celiac cells (6).

#### Combined IF and fluorescence in situ hybridization by RNAScope

Fluorescence in situ hybridization was performed on FFPE TMAs comprised of paired diagnostic and follow-up duodenal biopsies from CeD patients using RNAScope (77). Follow-up biopsies were obtained while patients were on a gluten-free diet (GFD) and showed reversion to normal or near normal histology. We used RNAScope Multiplex Fluorescent Assay with probes (Advanced Cell Diagnostics-ACD) against Epithelial Cell Adhesion Molecule (EpCAM) (#3120388), IFNG (#310508), CD3 (#426621), Granzyme B (#445978), TCR [illegible] chain (#433658), TCR [illegible] chain (#472438), CCL3 (#455331), CCL4 (#455341), CCL5 (#549171), CCL25 (#474381), CXCL9 (#440161), CXCL10 (#311851), and CXCL11 (#312701). Assays were either performed at ACD with images visualized using SlideView software or performed in our laboratory following the manufacturer's protocol (77). Sections were incubated with primary antibodies to CD3, EpCAM, Apolipoprotein A-IV (ApoA4) or collagen IV followed by fluorescence-conjugated secondary antibodies as described above. Following thorough washes in PBS, sections were visualized on the Stellaris 5 confocal microscope (Leica Microsystems Inc, Deerfield, IL 60015, United States). The total number of positive punctate dots representing RNA transcripts was counted in ten fields of view at 40x magnification on CD3+ cells and EpCAM+ enterocytes within ApoA4+ villous epithelium using ImageJ software version 1.53k. Negative control probe-processed sections were used as background signal and the average/mean quantified.

### Image Quantification and Statistics

Whole biopsy images were analyzed by tiling 20x images together using the EVOS Fl Auto 2 microscope or Stellaris confocal microscope. Regions of the villous and crypt epithelium were then selected based on ApoA4 and Ki67 stain, respectively. Quantification of antigens expressed on each part of the epithelium was performed using QuPath or ImageJ by calculating the mean fluorescence intensity per epithelial area or by measuring mean fluorescence intensity of antigen per CD3 cell or as a percentage of CD3 cells with bright or dim/absent staining. For RNAScope and PLA, the total number of spot/dots were counted in villous epithelium in ten fields of view at 40x magnification and results expressed either as average/mean of total number of RNA transcripts or as a percentage of PLA spots. Data are expressed as mean ± SEM. Statistical analyses were performed using GraphPad Prism software. The Mann-Whitney U test was used to make statistical comparisons between two unpaired groups while the Wilcoxon test was used for paired groups. A P value of less than or equal to 0.05 was considered statistically significant.

### Supplementary References


1. R. Elmentaite *et al.*, Cells of the human intestinal tract mapped across space and time. *Nature* **597**, 250-255 (2021).
2. P. L. Germain, A. Lun, C. Garcia Meixide, W. Macnair, M. D. Robinson, Doublet identification in single-cell sequencing data using scDblFinder. *F1000Res* **10**, 979 (2021).
3. I. Korsunsky *et al.*, Fast, sensitive and accurate integration of single-cell data with Harmony. *Nat Methods* **16**, 1289-1296 (2019).
4. C. Dominguez Conde *et al.*, Cross-tissue immune cell analysis reveals tissue-specific features in humans. *Science* **376**, eabl5197 (2022).
5. M. S. B. Raredon *et al.*, Comprehensive visualization of cell-cell interactions in single-cell and spatial transcriptomics with NICHES. *Bioinformatics* **39**, btac775 (2023).
6. D. Turei *et al.*, Integrated intra- and intercellular signaling knowledge for multicellular omics analysis. *Mol Syst Biol* **17**, e9923 (2021).

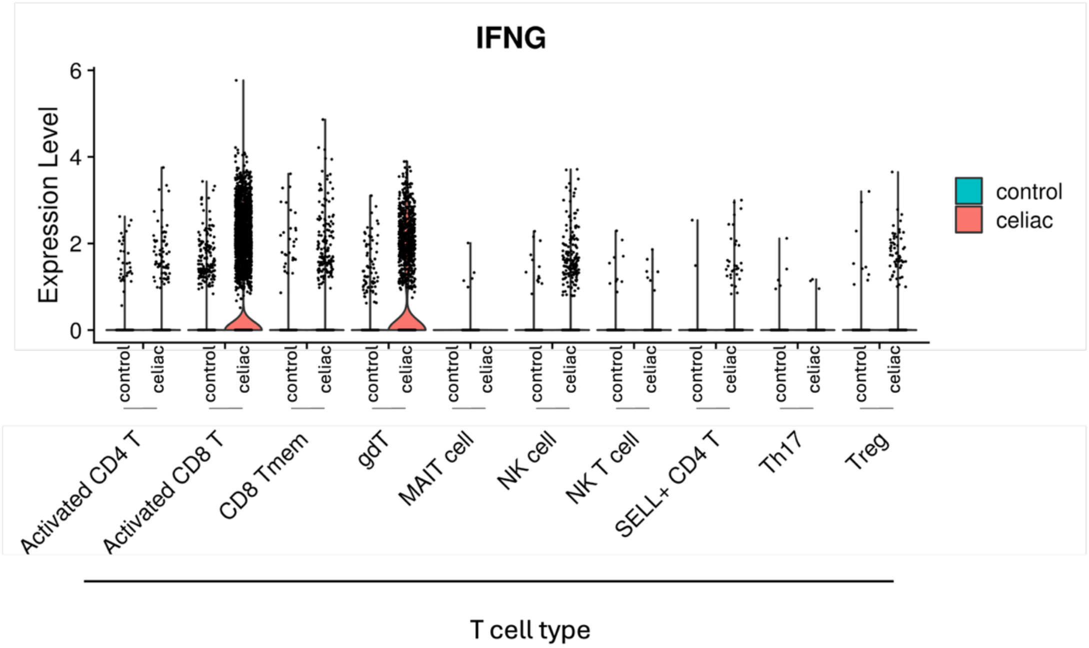


**Fig. S1. Single cell analysis of IFNG expression.** Violin plot of IFNG expression in T and NK cell subsets in control (blue) and celiac (pink) groups were analyzed using the 10x genomics scRNA-seq platform.

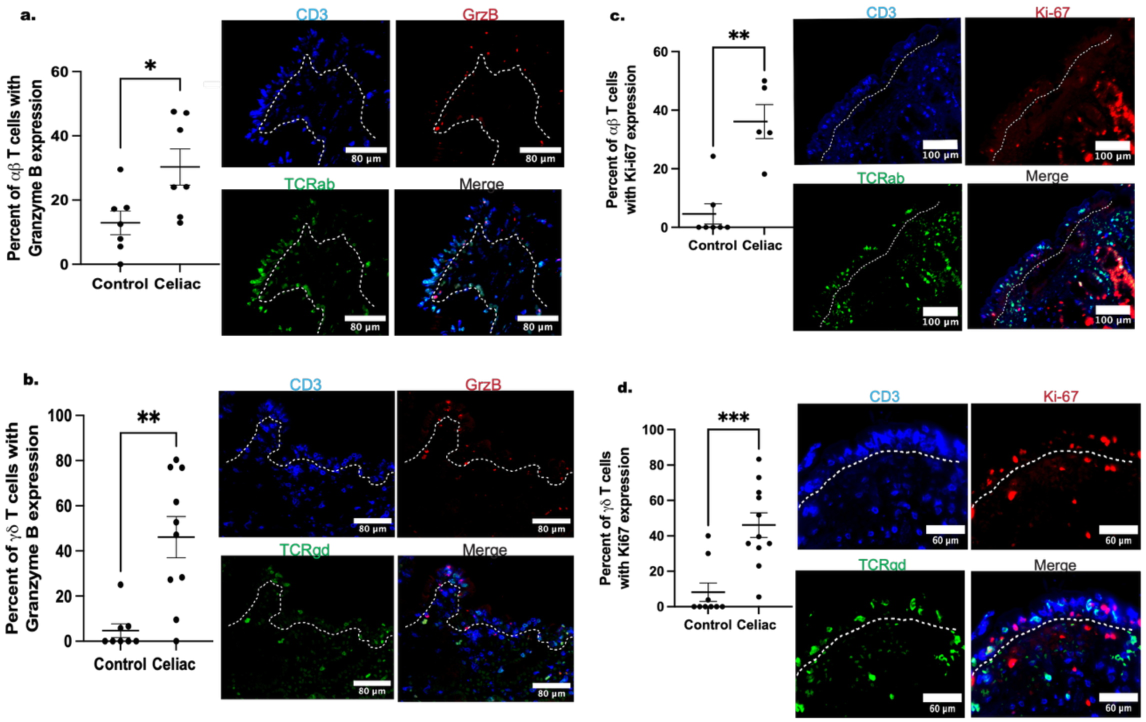


**Fig. S2. Expression of CTL markers within the villous epithelial compartment. (a.)** Representative IF images and quantification of Granzyme B expressing TCR αβ cells. The percentages of TCR αβ in the villous epithelium positive for Granzyme B are shown **(b).** Representative IF images and quantification of Granzyme B expressing TCR αβ cells. The percentages of TCR γδ in the villous epithelium positive for Granzyme B are shown. Ki-67 protein expression by IF in **(c.)** TCR αβ and **(d.)** TCR γδ T cell. Significance determined by t-test. *P ≤ 0.05; **P ≤ 0.01; ****P ≤ 0.001.

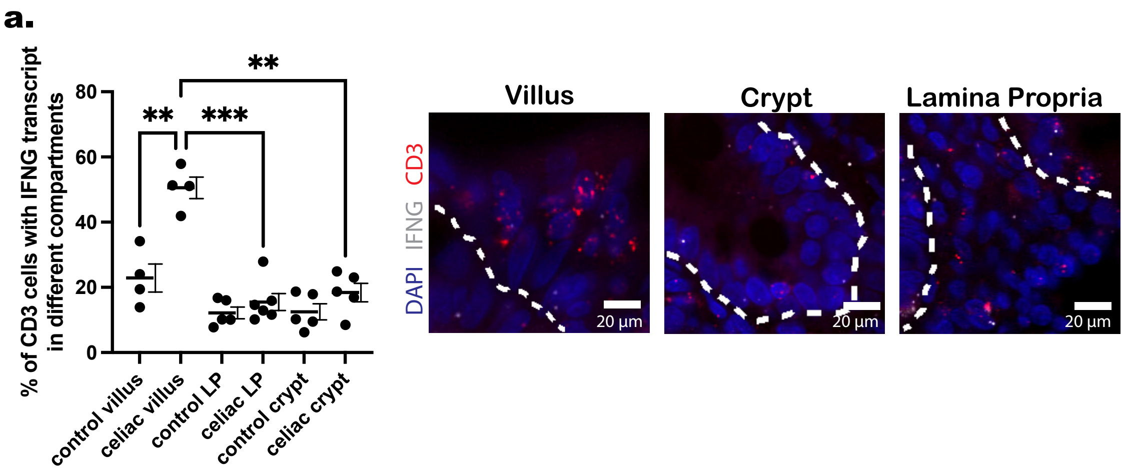


**Fig. S3. Assessment of IFNG transcript in celiac and control biopsies in different tissue compartments.** Quantification of the percentage of CD3 T cells with *IFNG* transcript in the villus, crypt, and lamina propria of celiac and control biopsies (N = 4). Significance determined by t-test. *P ≤ 0.05; **P ≤ 0.01; ****P ≤ 0.001

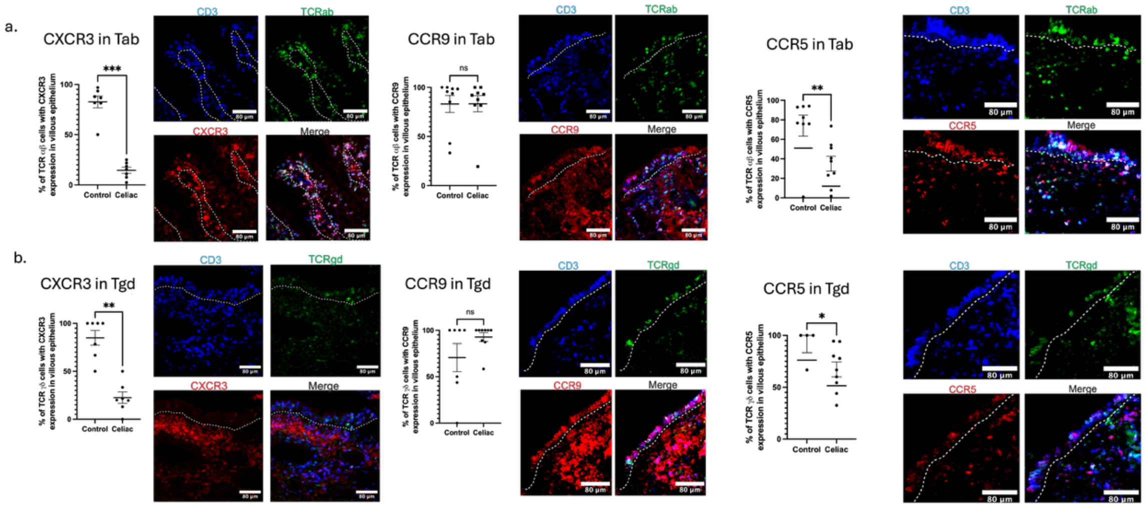


**Fig. S4. Chemokine receptor expression of TCRαβ and γδ populations found in the villi.** The percentage of **(a.)** αβ and **(b.)** γδ CD3 T cells that express chemokine receptors CXCR3, CCR5, and CCR9 in the villous. Each point represents one patient and the bars represent mean ± SEM. Significance was determined by t-test. *P ≤ 0.05; **P ≤ 0.01; ***P ≤ 0.001.

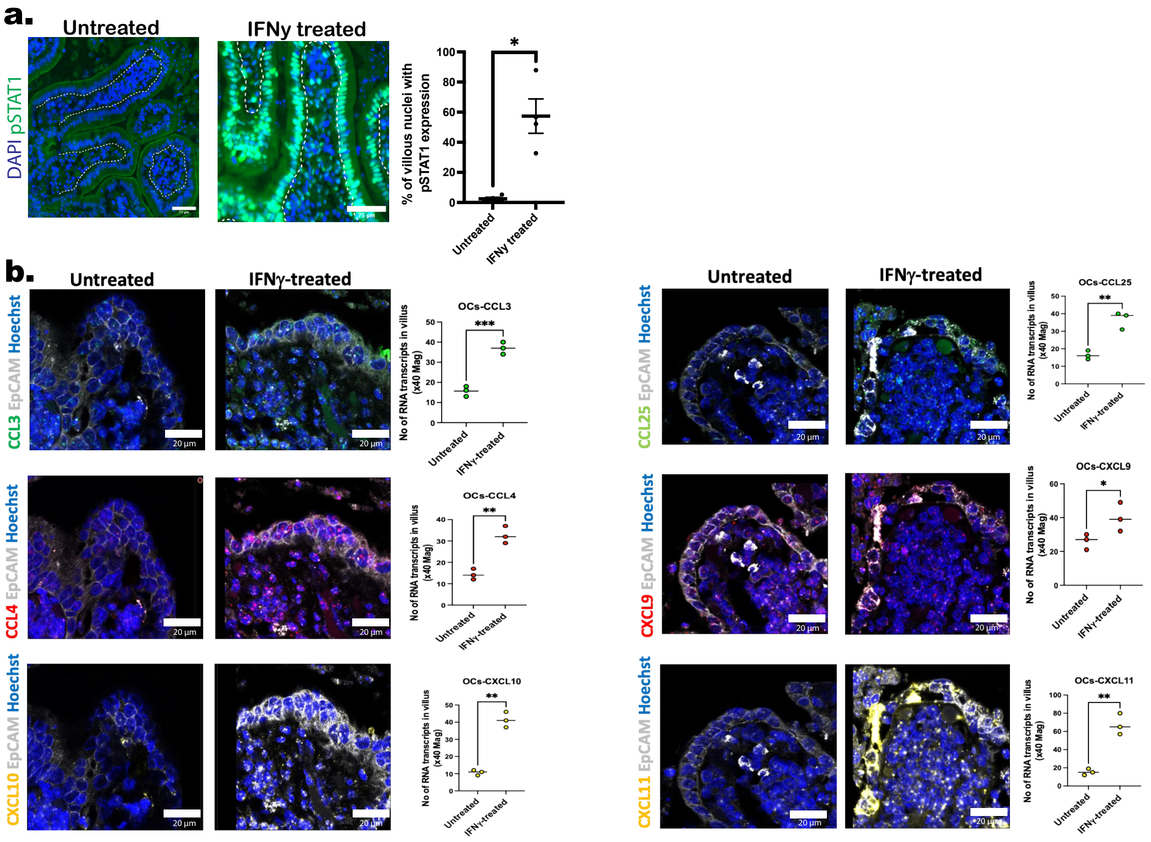


**Fig. S5. Induction of chemokine transcripts in duodenal mucosal organ cultures by IFNγ.** **(a.)** Human duodenal tissue from non-celiac patients were stimulated in vitro with IFNγ or left unstimulated. Cytokine activation was assessed by determining percentage of villous nuclei with pSTAT1 staining by IF. **(b.)** Human duodenal mucosal tissues were cultured in the presence or absence of IFNγ for 24 hours. Chemokine transcript expression of ligands CCL3, CCL4, CXCL9, CXCL10, CXCL11, and CCL25 was then assessed in the villous epithelium using RNAscope. Each point represents one patient and the bars represent mean ± SEM. Significance was determined by t-test. *P ≤ 0.05; **P ≤ 0.01; ***P ≤ 0.001.

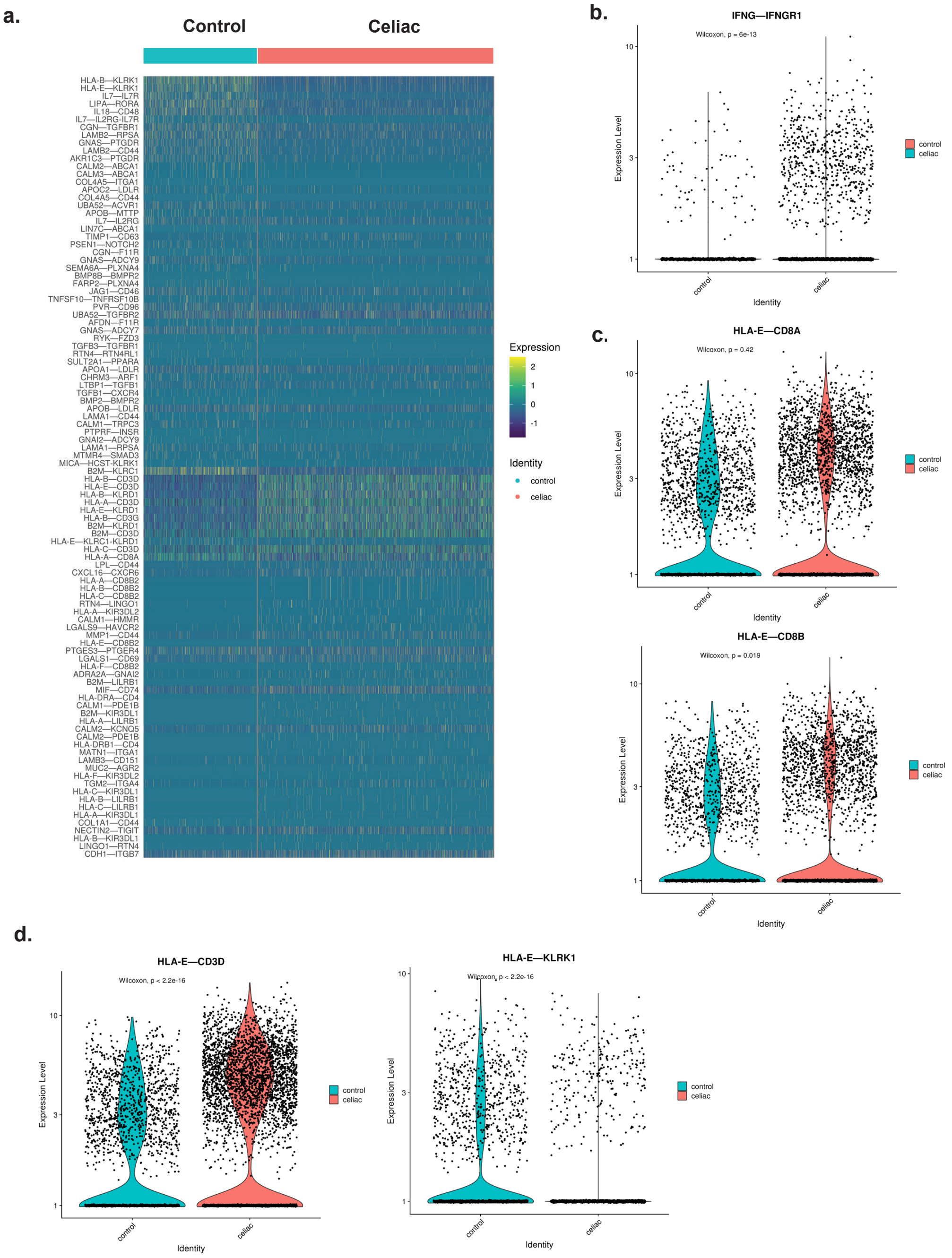


**Fig. S6. T cell/enterocyte interactions calculated from scRNA-seq.** (a). NICHES was used to find potential ligand receptor interactions from enterocytes to activated CD8T cells in our scRNA-seq dataset. The heatmap represents expression levels for the top 50 interactions from celiac and control enterocytes to activated CD8T cells. Each row is a ligand-receptor pair. Violin plots of the following selected ligand-receptor pairs found on enterocytes and activated CD8 T cells (b). IFNG-IFNG1, (c). HLA-E with TCR-related genes (CD3D, CD8A, CD8B). and (d). HLA-E and KLRC2. Significance determined by t-test. *P ≤ 0.05; **P ≤ 0.01; ****P ≤ 0.001.

**Table S1.** Patient demographics for TMA samples

Table S1. Patient demographics for TMA samples

**Supplementary Table 1. Patient demographics**

| Celiac patients (n=20)* | Age | Sex | Histology | Experiment | Figure | Controls (n= 35) | Age | Sex | Experiment | Figure |
|---|---|---|---|---|---|---|---|---|---|---|
| Patient 1 | 49 | F | Marsh grade 3 | IF | Fig. 2d,f; 4d,j; 6f; 7c,e,i | Control 1 | 17 | F | IF | Fig. 5b,e |
| Patient 2 | 5 | F | Marsh grade 3 | IF | Fig. 2d,f,e,g,h,I; 4b,d,j; 5b,e; 6d,f,h; 7g | Control 2 | 64 | M | IF | Fig. 5b |
| Patient 3 | 9 | M | Marsh grade 3 | IF & PLA | Fig. 2d,f,e,g,h,I; 4b,d,j; 5e; 6d,f,h; 7e,g | Control 3 | 58 | F | IF | Fig. 2d,f; 4d,j; 5b,e; 6d; 7c |
| Patient 4 | 6 | M | Marsh grade 3 | IF | Fig. 2d,f; 4d,j; 5e; 7e,g,i | Control 4 | 16 | M | IF | Fig. 5b,e |
| Patient 5 | 11 | M | Marsh grade 3 | IF | Fig. 3d,j; 5b; 6f; 7e,g | Control 5 | 9 | M | IF | Fig. 5b,e |
| Patient 6 | 61 | F | Marsh grade 3 | IF | Fig 5b; 7g,i | Control 6 | 56 | F | IF | Fig. 2d,f,e,g,h,i; 4d,j; 5b,e; 6f; 7c |
| Patient 7 | 23 | M | Marsh grade 3 | IF | Fig. 5b; 7c,g,i | Control 7 | 49 | F | IF & PLA | Fig. 2d,f; 4d,j; 5b; 6d; 7c |
| Patient 8 | 35 | F | Marsh grade 3 | IF | Fig 5b; 7g,i | Control 8 | 51 | F | IF | Fig. 2e,g,h,i; 5b; 6d |
| Patient 9 | 62 | F | Marsh grade 3 | IF | Fig. 2d,f; 4d; 5b,e; 6d,f,h; 7c,e,g | Control 9 | 63 | M | IF | Fig. 7i |
| Patient 10 | 17 | F | Marsh grade 3 | IF & PLA | Fig. 2d,f; 4b,d,j; 5b,e; 6d,f; 7c,e,g | Control 10 | 57 | M | IF | Fig. 6f; 7i |
| Patient 11 | 34 | M | Marsh grade 3 | IF | Fig. 5b,e | Control 11 | 53 | F | IF | Fig. 2d,f,e,g,h,i; 4d,j; 6d,f; 7c,e,g |
| Patient 12 | 24 | F | Marsh grade 3 | IF | Fig. 2d,f,e,g,h,I; 4d,j; 5b,e; 6d,h; 7c,g,i | Control 12 | 68 | F | IF | Fig. 6f; 7i |
| Patient 13 | 71 | M | Marsh grade 3 | IF | Fig. 2e,g,h,I; 4j; 5b,e; 6d,f,h; 7c,g | Control 13 | 62 | F | IF | Fig. 6f; 7i |

| | | | | | | | | | |
|---|---|---|---|---|---|---|---|---|---|
| Patient 14 | 64 | F | Marsh grade 3 | IF | Fig. 2d,f,e,l; 4d,j; 5b,e; 6d,f; 7c,e,g | Control 14 | 34 | F | IF | Fig. 6f; 7i |
| Patient 15 | 15 | F | Marsh grade 3 | IF | Fig. 2i; 4j; 5b,e; 6h; 7c,e,g,i | Control 15 | 50 | F | IF | Fig. 6f |
| Patient 16 | 12 | M | Marsh grade 3 | IF | Fig. 5b; 6d,f | Control 16 | 22 | F | IF | Fig. 6f; 7i |
| Patient 17 | 5 | M | Marsh grade 3 | IF | Fig. 2d,f,e,g,h,I; 4d,j; 5b,e; 6f; 7c,e,g,i | Control 17 | 69 | F | IF | Fig. 2d,e,f,g,h,i; 4d,j; 6f; 7c,e,g,i |
| Patient 18 | 38 | M | Marsh grade 3 | IF | Fig. 2e,g,h; Fig. 4b,d,j; 5b,e; 6d; 7c,e,g,i | Control 18 | 68 | F | IF | Fig. 7i |
| Patient 19 | 24 | F | Marsh grade 3 | IF | Fig. 6f | Control 19 | 40 | F | IF | |
| Patient 20 | 19 | F | Marsh grade 3 | IF | Fig. 7i | Control 20 | 61 | F | IF | Fig. 2e,g,h,i; 5e; 6d,h; 7c,e,g |
| | | | | | | Control 21 | 72 | F | IF | Fig. 4b,d; 5e; 6h; 7c,e |
| | | | | | | Control 22 | 51 | F | IF | Fig. 4b,d; 5e; 6h; 7c,e,g |
| | | | | | | Control 23 | 82 | F | IF | Fig. 2e; 4b,d; 5e; 6h; 7c,e,g |
| | | | | | | Control 24 | 70 | F | IF | Fig. 4b,d; 5e; 6d; 7c,e,g |
| | | | | | | Control 25 | 16 | M | IF | Fig. 5b,e; 7g |
| | | | | | | Control 26 | 3 | F | IF | Fig. 2d,f; 4d,j; 5b,e; 6h; 7c,e,g |
| | | | | | | Control 27 | 58 | M | IF | Fig. 2e,g,h,i; Fig. 4b,j; 5b,e; 6d,h; 7g |
| | | | | | | Control 28 | 51 | M | IF & PLA | Fig. 2d,f; 4b,d,j; 5b; 6h; 7c,e,g |
| | | | | | | Control 29 | 69 | F | IF | Fig. 2e; 4j; 6d,f; 7c,e,g |

| | | | | |
|---|---|---|---|---|
| Control 30 | 67 | M | IF | Fig. 2d,f; Fig. 4b,d,j; 5e; 7g |
| Control 31 | 11 | F | IF | Fig. 2d,f; 4d,j; 5b,e; 7e,g,i |
| Control 32 | 8 | F | IF | Fig. 5b 7g |
| Control 33 | 10 | M | IF | Fig. 5b; 7e,g |

**Table S2.** Patient demographics for TMA samples

**Supplementary Table 2. Patient demographics for TMA samples**

**TMA 1**

| Celiac patients (n = 6) | Age | Sex | Histology | Experiment | Controls (n= 6) | Age | Sex | Experiment |
|---|---|---|---|---|---|---|---|---|
| Patient 1 | 49 | F | Marsh grade 3 | FISH & PLA | Control 1 | 20 | M | FISH & PLA |
| Patient 2 | 6 | M | Marsh grade 3 | FISH & PLA | Control 2 | 13 | F | FISH & PLA |
| Patient 3 | 31 | F | Marsh grade 3 | FISH & PLA | Control 3 | 6 | F | FISH & PLA |
| Patient 4 | 15 | M | Marsh grade 3 | FISH & PLA | Control 4 | 59 | F | FISH & PLA |
| Patient 5 | 3 | F | Marsh grade 3 | FISH & PLA | Control 5 | 45 | F | FISH & PLA |
| Patient 6 | 7 | M | Marsh grade 3 | FISH & PLA | Control 6 | 10 | M | FISH & PLA |

**TMA 2**

| Celiac patients (n = 6) | Age | Sex | Histology | Experiment | Controls (n= 6) | Age | Sex | Experiment |
|---|---|---|---|---|---|---|---|---|
| Patient 1 | 33 | F | Marsh grade 3 | FISH & PLA | Control 1 | 13 | F | FISH & PLA |
| Patient 2 | 17 | F | Marsh grade 3 | FISH & PLA | Control 2 | 55 | F | FISH & PLA |
| Patient 3 | 28 | F | Marsh grade 3 | FISH & PLA | Control 3 | 53 | M | FISH & PLA |
| Patient 4 | 46 | M | Marsh grade 3 | FISH & PLA | Control 4 | 69 | F | FISH & PLA |
| Patient 5 | 30 | F | Marsh grade 3 | FISH & PLA | Control 5 | 25 | F | FISH & PLA |
| Patient 6 | 4 | F | Marsh grade 3 | FISH & PLA | Control 6 | 11 | F | FISH & PLA |

**TMA 3: initial diagnostic and f/u biopsies on GFD**

| Active CeD (n = 5) | Age | Sex | Histology | Experiment | GFD (n= 4) | Age | Sex | Experiment |
|---|---|---|---|---|---|---|---|---|
| Patient 1 | 69 | F | Marsh grade 3 | IF & FISH | Patient 1 | 69 | F | IF & FISH |
| Patient 2 | 53 | F | Marsh grade 3 | IF & FISH | Patient 2 | 60 | F | IF & FISH |
| Patient 3 | 52 | M | Marsh grade 3 | IF & FISH | Patient 3 | 53 | M | IF & FISH |
| Patient 4 | 12 | M | Marsh grade 3 | IF & FISH | Patient 4 | 66 | M | IF & FISH |
| Patient 5 | 16 | F | Marsh grade 3 | IF & FISH | | | | |

**Table S3.** List of Antibodies used for this study.

| Supplementary Table 3. List of Antibodies used for this study | | | | |
|---|---|---|---|---|
| **Antigen** | **Clone** | **Catalog #** | **Source** | **Dilution** |
| Apo A4 | G-8 | Sc 374543 | Santa cruz | 1:100 |
| B2 microglobulin | D8P1H | 59035 | CST | 1:200 |
| CCR5 | polyclonal | 17476-1 | Proteintech | 1:100 |
| CCR9 | polyclonal | PA5-33405 | Thermo fisher | 1:200 |
| CD56 | 123c3 | Ma5 16446 | Thermo fisher | 1:100 |
| CD45RO | Uchl1 | 304202 | Biolegend | 1:100 |
| CD3 | CD3-12 | ab11089 | Abcam | 1:100 |
| CD8a | C8/144B | 70306S | CST | 1:100 |
| Collagen IV | polyclonal | NBP1-26549 | Novus bio | 1:500 |
| CXCR3 | EPR25373-32 | ab288437 | Abcam | 1:100 |
| DAP12 | G-5 | Sc 133174 | Santa Cruz | 1:100 |
| DAP12 | E8P9U | 46088 | CST | 1:100 |
| EpCAM | monoclonal | MA5-29246 | Novus Bio | 1:200 |
| E-cadherin | 24E10 | CST 3199 | CST | 1:100 |
| Granzyme B | D6E9W | 46890S | CST | 1:100 |
| HLA-B | polyclonal | PA5-35345 | Thermo Fisher | 1:200 |
| HLA-DR | TAL 1B5 | Ab20181 | Abcam | 1:200 |
| HLA-E | polyclonal | 27411-1 | Proteintech | 1:50 |
| HLA-E | 1A4G3 | 66530-1-Ig | Proteintech | 1:400 |
| Ki67 | SP6 | Ab16667 | Abcam | 1:200 |
| NKG2c | polyclonal | GTX81130 | GeneTex | 1:100 |
| NKG2c | polyclonal | LS-B13756 | LSBio | 1:200 |
| Nur77 | NR4A1 | PA5-18573 | Thermo Fisher | 1:100 |
| Pan cytokeratin | AE-1/A-3 | Nbp2-29429 | Novus bio | 1:100 |
| Phospho-STAT1 | 58D6 | CST 9167 | CST | 1:200 |
| TCR $\alpha\beta$ | polyclonal | 11890-1-AP | Proteintech | 1:100 |
| TCR $\alpha\beta$ | H-1 | Sc-515719 | Santa Cruz | 1:33 |
| TCR $\gamma\delta$ | H-41 | Sc-100289 | Santa Cruz | 1:33 |
| TCR $\gamma\delta$ | monoclonal | 5570T | CST | 1:100 |

**Table S4.** Chemokine ligand mRNA spots per field in organ culture.

**Supplementary Table 4. Chemokine ligand mRNA spots per field in organ culture**

| Chemokine ligand | Control | Celiac | P value |
|---|---|---|---|
| CCL3 | 15.6 ± 0.1 | 37.1 ± 0.2 | *** |
| CCL4 | 14.3 ± 0.1 | 32.6 ± 0.2 | ** |
| CXCL9 | 31.1 ± 0.7 | 40.2 ± 0.3 | * |
| CXCL10 | 10.6 ± 0.3 | 41.3 ± 0.2 | ** |
| CXCL11 | 15.3 ± 0.2 | 67.3 ± 0.4 | ** |
| CCL25 | 16.3 ± 0.2 | 36.6 ± 0.4 | ** |

Table shows the average number of mRNA hybridized spots in the villous per field in organ cultures of human normal duodenal tissues stimulated with IFN-y or left unstimulated. Data presented (mRNA spots per 40x magnification) are means ± S.E.M., n = 3 individual biopsies each with ten images. **p < 0.001 by Mann-Whitney test.
